# Transcriptome Analysis of Distinct Cold Tolerance Strategies in the Rubber Tree (*Hevea brasiliensis*)

**DOI:** 10.1101/395590

**Authors:** Camila Campos Mantello, Lucas Boatwright, Carla Cristina da Silva, Erivaldo Jose Scaloppi, Paulo de Souza Gonçalves, W. Brad Barbazuk, Anete Pereira de Souza

## Abstract

Natural rubber is an indispensable commodity used in approximately 40,000 products and is fundamental to the tire industry. Among the species that produce latex, the rubber tree [*Hevea brasiliensis* (Willd. ex Adr. de Juss.) Muell-Arg.], a species native to the Amazon rainforest, is the major producer of latex used worldwide. The Amazon Basin presents optimal conditions for rubber tree growth, but the occurrence of South American leaf blight, which is caused by the fungus *Microcyclus ulei* (P. Henn) v. Arx, limits rubber tree production. Currently, rubber tree plantations are located in scape regions that exhibit suboptimal conditions such as high winds and cold temperatures. Rubber tree breeding programs aim to identify clones that are adapted to these stress conditions. However, rubber tree breeding is time-consuming, taking more than 20 years to develop a new variety. It is also expensive and requires large field areas. Thus, genetic studies could optimize field evaluations, thereby reducing the time and area required for these experiments. Transcriptome sequencing using next-generation sequencing (RNA-seq) is a powerful tool to identify a full set of transcripts and for evaluating gene expression in model and non-model species. In this study, we constructed a comprehensive transcriptome to evaluate the cold response strategies of the RRIM600 (cold-resistant) and GT1 (cold-tolerant) genotypes. Furthermore, we identified putative microsatellite (SSR) and single-nucleotide polymorphism (SNP) markers. Alternative splicing, which is an important mechanism for plant adaptation under abiotic stress, was further identified, providing an important database for further studies of cold tolerance.

## 1 Introduction

Cold stress, which can be classified as chilling (0 to 15°C) and/or freezing (< 0°C) temperatures, affects plant growth and development, limiting spatial distribution and yields (Chinnusamy et al., 2007; Yang et al., 2015). Furthermore, cold stress prevents plants from achieving their full genetic potential, inhibiting metabolic reactions, reducing photosynthetic capacity and altering membrane permeability (Chinnusamy et al., 2007; Sevillano et al., 2009).

Temperate plants can generally achieve cold acclimation and acquire tolerance to extracellular ice formation in their vegetative tissues. However, tropical crops, such as maize and rice, lack cold acclimation ability and are sensitive to chilling (Chinnusamy et al., 2007). Furthermore, varieties from the same species can exhibit different levels of cold tolerance (Liu et al., 2012; Zhang et al., 2012). Hence, determining the gene expression profile under cold stress could help to elucidate the mechanism of cold acclimation in plants and can be effective for selecting candidate genes. Moreover, candidate genes can be targeted to identify genetic variation and develop molecular markers.

*Hevea brasiliensis* [(Willd. ex Adr. de Juss.) Muell-Arg], commonly known as the rubber tree, is a perennial tree crop native to the Amazon rainforest. The species, belonging to the Euphorbiaceae family, is monoecious, undergoes cross-pollination and has a chromosome number of 2n = 2x = 36. Among the 2,500 species that produce natural rubber (cis-1,4-polyisoprene), *H. brasiliensis* is the only species that produces high-quality rubber in commercially viable quantities, accounting for more than 98% of total worldwide production (Pootakham et al., 2017).

Natural rubber is one of the most important raw materials for many industries and cannot be replaced by synthetic alternatives due to its unique properties, such as flexibility, elasticity and abrasion resistance (Sakdapipanich, 2007). Natural rubber is an essential commodity for the tire industry and for the manufacture of more than 40,000 products.

Although the Amazon basin offers a suitable climate for this crop, Southeast Asia is the major producer of rubber, being responsible for 92% of worldwide production. South America is responsible for only 2% of worldwide rubber production, due to the occurrence of the fungus *Microcyclus ulei* (P. Henn) v. Arx, which causes South American leaf blight (SALB). SALB was responsible for devastating plantations in northern Brazil in the 1930s and remains a permanent threat to the rubber industry (Pushparajah, 2001). To date, the rubber tree plantations in Southeast Asia have not been affected by SALB, but other native pathogenic fungi are threats to rubber production. The two major fungal pathogens in Southeast Asia (*Phytophthora* and *Corynespora*) cause leaf fall and, consequently, significant losses of natural rubber yields (Pootakham et al., 2017). Due to the occurrence of diseases, plantations have been expanded to sub-optimal areas of some countries, such as northeastern India, the highlands and coastal areas of Vietnam, southern China and the southern plateau of Brazil (Priyadarshan et al., 2005). These areas are characterized by new stressful conditions, such as cold and dry periods.

The exposure of rubber trees to low temperatures can cause leaf necrosis, affecting tree development and latex production (Priyadarshan et al., 2005; Mai et al., 2010). In addition, low temperatures are responsible for halting latex production for 1–3 months per year (Rao et al., 1998; Jacob et al., 1999).

In recent years, there has been an exponential increase in genomic data acquisition for the rubber tree, including transcriptome profiles (Mantello et al., 2014; Salgado et al., 2014; Li et al., 2016), linkage maps (Souza et al., 2013; Pootakham et al., 2015; Shearman et al., 2015) and, more recently, a genome assembly (Tang et al., 2016; Pootakham et al., 2017). Despite the importance of the cold acclimation response for rubber tree breeding programs, no studies conducted to date have focused on identifying the set of genes involved in the response to chilling stress tolerance.

To understand this mechanism, we conducted a chilling stress experiment (10°C) with the clones GT1 and RRIM600, which exhibit high yields and are recommended for planting in escape areas. The clone RRIM600 is a cold resistant clone that stops growing under cold stress, while GT1 is chilling tolerant, showing little leaf damage and continuing to grow under chilling temperatures (Mai et al., 2010). RNA sequencing was performed with the aim of constructing a comprehensive transcriptome and investigating the differentially expressed genes (DEGs) involved in different cold acclimation strategies. In addition, the comprehensive transcriptome was searched for putative molecular markers (single-nucleotide polymorphisms (SNPs) and microsatellites) and to detect alternative splicing (AS) events. Alternative splicing is an important mechanism responsible for generating transcriptome diversity, where resulting splicing variants may perform a variety of functions, such as increase cold resistance (Chinnusamy et al., 2007; Tack et al., 2014).

## 2 Material and Methods

### 2.1 Plant Materials and Cold Stress Treatment

Plantlets of the rubber tree clones RRIM600 and GT1 at 6 months of age were provided by Centro de Seringueira, Votuporanga, Sao Paulo, Brazil. The clones were represented by 3 biological replicates each.

The plants were transferred to a growth chamber set to 28°C with a 12 h photoperiod and were watered every 2 days for 10 days. After 10 days, the plantlets were exposed to chilling stress at 10°C for 24 h. Leaf tissues were sampled at 0 h (control), 90 minutes, 12 h and 24 h after cold exposure. The samples were immediately frozen on dry ice and stored at −80°C until use.

### 2.2 RNA Extraction and cDNA Library Construction and Sequencing

Total RNA was extracted using a modified lithium chloride protocol (Oliveira et al., 2015). RNA integrity and quantity were assessed using an Agilent 2100 Bioanalyzer (Agilent Technologies, Palo Alto, CA). Approximately 3 µg of total RNA was employed to construct cDNA libraries using the TruSeq RNA Sample Preparation Kit (Illumina Inc., San Diego, CA, USA). Index codes were added to each sample, and the cDNA libraries were prepared following the manufacturer’s recommendations. In total, 24 cDNA libraries (3 replicates of each genotype for each time series) were prepared. Library quality was evaluated with a 2100 Bioanalyzer (Agilent Technologies, Palo Alto, CA), and the libraries were quantified via qPCR (Illumina protocol SY-930-10-10). The 24 samples were randomly pooled (4 samples per pool) and clustered using the TruSeq PE Cluster Kit on the cBot platform (Illumina Inc., San Diego, CA, USA). Subsequently, the cDNA libraries were sequenced using an Illumina Genome Analyzer IIx with the TruSeq SBS 36-Cycle Kit (Illumina, San Diego, CA, USA) for 72 bp paired-end reads.

### 2.3 Data Filtering

The raw data generated via Illumina sequencing in the BCL format were converted to the qSeq format using Off-Line Basecaller v.1.9.4 (OLB) software. We further converted the qSeq files into FastQ files using a custom script. Therefore, the raw reads were split by the corresponding barcodes, and the barcode regions were trimmed using the Fastx-Toolkit (fastx_barcode_splitter.pl) (http://hannonlab.cshl.edu/fastx_toolkit/index.html).

Filtering for high-quality (HQ) reads was performed using NGS QC Toolkit 2.3 (Patel and Jain, 2012), considering only reads with a Phred quality score ≥ 20 and a cut-off value of 70% of read length. All reads were deposited in the NCBI Short Read Archive (SRA) under accession number SRP155829.

### 2.4 Comprehensive Transcriptome Assembly

Initially, the reads generated from the 24 samples were combined with bark reads (Mantello et al., 2014) and mapped back onto the rubber tree genome (Tang et al., 2016) (accession number: LVXX01000000) with the HISAT2 aligner (Kim et al., 2015). The alignment results were coordinate-sorted with SAMtools (Li et al., 2009) and used to perform Trinity genome-guided assembly. Furthermore, the scaffolds obtained from the rubber tree genome were employed for an *ab initio* genome annotation with MakerP (Campbell et al., 2014) (Supplementary Material 1) due to the lack of public genome annotation. This annotation provided an additional dataset of predicted transcripts.

The transcripts obtained in the genome-guided and *ab initio* genome annotations were combined with non-redundant *H. brasiliensis* ESTs from NCBI (as for Ago 2016) and used as a dataset for alignment and assembly against the rubber tree genome (Tang et al., 2016), using the PASA v2.0 pipeline (Haas et al., 2003) with the following parameters: --ALT_SPLICE --ALIGNER blat,gmap and MAXIMUM_INTRON_LENGTH=“50000”. PASA modeled complete and partial gene structures based on splice-aware alignment to a reference genome, detecting unique assemblies, collapsing redundant models and identifying AS events.

The transcripts obtained using PASA were filtered according to the following criteria: (1) minimum length of 500 bp; (2) transcript prediction evidence, excluding transcripts that were exclusively predicted in the genome *ab initio* annotation; and (3) trimming of transcripts with high identity to non-plant sequences.

The filtered transcripts were clustered based on genome mapping location and according to gene structures. This final dataset was considered the comprehensive transcriptome for further analysis.

### 2.5 Functional Annotation

The Trinonate v2.0.1 pipeline (https://trinotate.github.io/) was employed to annotate the transcriptome. Briefly, Transdecoder (https://github.com/TransDecoder/TransDecoder/wiki) was used to predict open reading frames (ORFs) with a minimum of 100 amino acids. Transdecoder can predict multiple ORFs in the same transcript; however, if the predicted ORFs overlap, the program maintains the longest ORF. If multiple non-overlapping ORFs are predicted in the same transcript, all are retained in the annotation. Translated ORFs and untranslated transcripts are searched against the SwissProt/UniProt database using BLASTX and BLASTP, respectively. In addition, these transcripts were associated with Gene Ontology (GO) (Harris et al., 2004) and Kyoto Encyclopedia Gene and Genomes (KEGG) (Kanehisa and Goto, 2000) database information. The Transdecoder-predicted proteins were also searched for protein domain homology in the Pfam database using the HMMER 3.1 tool hmmscan (hmmer.org.). All the annotations were filtered with an e-value of 1e-5 and placed into a tab-delimited file.

### 2.6 Differential Gene Expression

Reads from each library were aligned to the reference transcriptome with Bowtie2 v.2.2.6 (Langmead and Salzberg, 2012), and the estimation of gene transcript abundance was performed with RSEM v.1.2.28 (Li and Dewey, 2011) using a Trinity accessory script (align_and_estimate_abundance.pl). The differential gene expression analysis was performed with limma-voom (Law et al., 2014), which estimates precision weights based upon an expression mean-variance trend to facilitate Bayesian-moderated, weighted t-stastitics (Soneson and Delorenzi, 2013), with at least 10 counts per million (CPM) in at least 3 samples. Three biological replicates for each condition were provided for this analysis. We considered a gene to be differentially expressed using a false discovery rate (FDR) cut-off ≤ 0.05. The pairwise comparison was performed between RRIM600 and GT1 for each time-series of the cold treatment: RRIM600 0h x GT1 0h, RRIM600 90min x GT1 90min, RRIM600 12h x GT1 12h, RRIM600 24h x GT1 24h.

### 2.7 Gene Ontology Enrichment

The DEGs identified previously were subjected to GO enrichment analysis using GOseq with a FDR cut-off ≤ 0.05 (Young et al., 2010). The enriched terms were submitted to REVIGO (Supek et al., 2011) with a medium similarity allowed (0.7) to summarize the enriched terms.

### 2.8 Putative Molecular Marker Identification

Putative Microsatellites (SSRs) were identified using the MISA (MIcroSAtellite) script (http://pgrc.ipk-gatersleben.de/misa/). SSR regions were defined as containing at least a six motif repetition for dinucleotides and 5 for tri-, tetra-, penta-and hexanucleotides.

The identification of putative SNPs was performed for each genotype. The reads obtained in this study were mapped against the reference transcriptome with bwa-mem (Li and Durbin, 2009) following the default parameters. SAM files were converted into BAM files using SAMtools. Additionally, we used SAMtools to sort mapped reads and remove unmapped reads. PCR duplicates were removed with Picard (http://broadinstitute.github.io/picard). The software Freebayes (Garrison and Marth, 2012) was used to call variants in each processed BAM file with the following parameters: --min-alternate-count 5 --min-mapping-quality 30 --min-base-quality 20. VCFtools (Danecek et al., 2011) was used to select biallelic SNPs, remove Indels, and perform filtering with a minimum genotype quality of 20, minimum depth of 10 reads and SNP and mapping quality of 20.

### 2.9 Quantitative RT-PCR (qRT-PCR) Validation

To validate the DEG analysis, a total of 20 genes were selected. Primer pairs used in the qRT-PCR analyses were designed using Primer3 Plus software (http://www.bioinformatics.nl/cgi-bin/primer3plus/primer3plus.cgi) using the qPCR parameters. cDNA synthesis was performed with the Quantitect Reverse Transcription kit (Qiagen Inc., Chatsworth, CA, USA) using 500 ng of total RNA. The cDNAs were then diluted 1:5, and 2 µl from each sample aliquoted for qPCR. The qPCR assays were carried out with iTaq Universal SYBR® Green Supermix (Bio-Rad Laboratories Inc., Hercules, CA, USA), following the manufacturer’s instructions, and 3 µM primer mixture. The qPCR assays were performed using the CFX384 Real-Time PCR Detection System, with the following cycling conditions: 95°C for 10 min, followed by 40 cycles of 95°C at 30 s and 60°C at 1 min.

All qPCR experiments were performed using three technical and three biological replicates, with the exception of RRIM 600 at 0 h and 12 h of treatment, for which only two biological replicates were included due to the lack of RNA for one of the biological replicates. The DEAD box RNA helicase (RH2b) and mitosis protein (YSL8) genes were used as internal controls. To confirm the presence of a single amplicon of the PCR product, melting curve analysis was performed with temperatures ranging from 65°C to 95°C in increments of 0.5°C. The Cq values and baseline were determined with CFX Manager 2.1 software (Bio-Rad Laboratories, Inc., USA). The primers used in this study are described in Supplementary Table 1.

### 2.10 Alternative Splicing Identification

The filtered transcripts and AS events defined by PASA were processed using an in-house pipeline. This pipeline identifies and re-classifies AS events simultaneously encompassing alternative 5’ and 3’ splice sites (Chamala et al., 2015). Furthermore, a minimum of 10 reads mapped at the splice junction was set as the threshold for considering an AS event.

## 3 Results

### 3.1 Sequencing and Transcriptome Assembly

In the present study, we sequenced leaf tissue from the RRIM600 and GT1 genotypes. Twenty-four cDNA libraries were sequenced on the Illumina GAIIx platform, which resulted in a total of 529,339,330 M paired-end (PE) reads for the RRIM600 genotype and 632,887,764 M PE reads for GT1. After removing low-quality reads, the cDNA libraries derived from RRIM600 yielded 432,005,062 M (81.6%) HQ PE reads, while GT1 yielded 501,609,042 M (79.2%) HQ PE reads (Table 1). When we summarized the total HQ PE reads from both genotypes, we obtained 933,614,104 M reads, which were employed to construct the reference transcriptome.

**Table 1.**
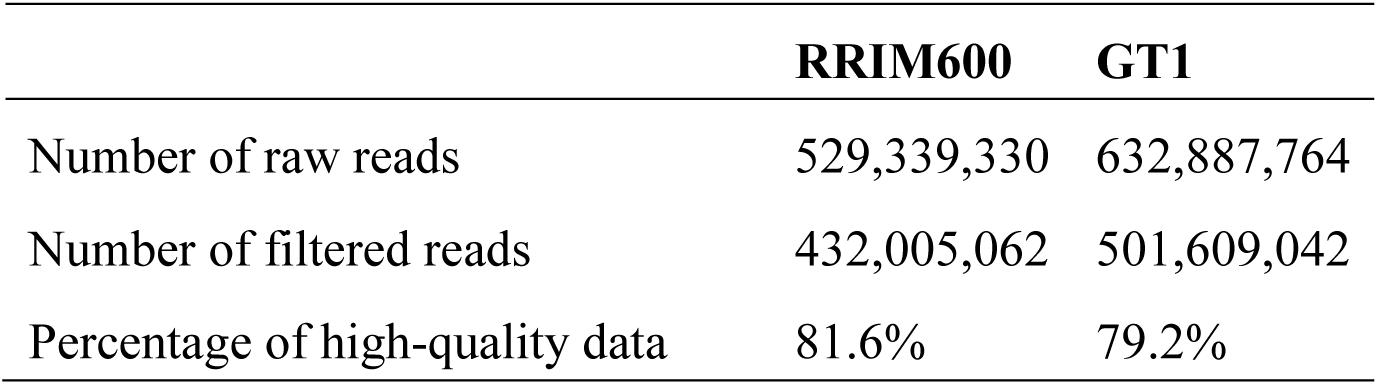
Summary of Illumina Sequencing.

To generate a comprehensive transcriptome, we used the PASA pipeline to update and maximize the recovery of gene structure and spliced isoforms. The 39,351 non-redundant ESTs from NCBI were combined with 335,212 transcripts obtained in the genome-guided assembly and the 114,718 filtered transcripts obtained from genome annotation (Supplementary Material 1) and then aligned against the rubber tree genome (Tang et al., 2016).

A total of 250,458 transcripts were obtained, which clustered with 162,278 genes. The N50 was 2,095 pb, and the GC content was 40.31%. These transcripts were filtered out according to their length (≤ 500 bp), their similarity to non-plant sequences via BLASTX and whether the genome annotation was the only evidence of the prediction. After filtering the transcripts obtained using PASA, a total of 104,738 transcripts were obtained and clustered with 49,304 genes (Table 2). Although a public annotation is not available, the total number of predicted genes in this study was similar to the prediction of Tang et al. (2016) for the rubber tree genome assembly (43,792 genes). The N50 of the reference transcriptome obtained in this study was 2,369 bp with a GC% content of 40.16% (Table 2). Of the total transcripts, 37,302 (35.6%) ranged in size from 1,000 bp to 1,999 bp, and 36,681 (35%) transcripts were longer than 2 kb. The 104,738 transcripts were considered a reference transcriptome and employed for further analysis.

**Table 2.** Summary statistics for the comprehensive transcriptome.

|  |  |
| --- | --- |
| Total number of contigs | 104,738 |
| Total number of genes | 49,304 |
| Total number of nucleotides | 196,309,369 |
| Min contig length (bp) | 500 |
| Max contig length (bp) | 22,333 |
| Mean contig Length (bp) | 1,874 |
| N50 | 2,369 |
| GC content | 40.16% |

### 3.2 Functional Annotation

The transcriptome was subjected to BLAST searches against the SwissProt/UniProt, Pfam, Gene Ontology and KEGG databases with an e-value cut-off of 1e-5. In total, 63,983 (61%) transcripts were annotated against the SwissProt/UniProt database using BLASTX. In addition, among the 94,166 proteins predicted with Transdecoder, 42.262 (44.9%) were annotated based upon BLASTP.

The Pfam annotation contained protein domains for 67,628 (71.8%) predicted proteins. The top 20 protein domains are presented in Figure 1. The most abundant type of protein domain was a protein kinase domain, which was found in 2,415 proteins. The protein kinase superfamily catalyzes the reversible transfer of the gamma-phosphate from ATP to amino acid side chains of proteins. These enzymes are involved in the response to many signals, including light, pathogen invasion, hormones, temperature stress, and nutrient deprivation. The activities of several plant metabolic and regulatory enzymes are also controlled by reversible phosphorylation (Stone and Walker, 1995). The next three most abundant gene families identified were the protein tyrosine kinase, leucine-rich repeat (LRR) N-terminal domain and NB-ARC domain families, with 1,055, 962 and 868 proteins, respectively (Figure 1). The protein tyrosine kinase family is responsible for signal transduction in plants in response to stress and developmental processes through modification of tyrosine residues in proteins (Shankar et al., 2015). Proteins containing the LRR N-terminal domain are involved in numerous functions, such as signal transduction, cell adhesion, DNA repair, disease resistance, apoptosis and the immune response. The NB-ARC domain is a functional ATPase, and its nucleotide-binding state regulates the activity of resistance (R) proteins. R proteins are involved in pathogen recognition, which activates the immune system in plants (van Ooijen et al., 2008).

**Figure 1.**
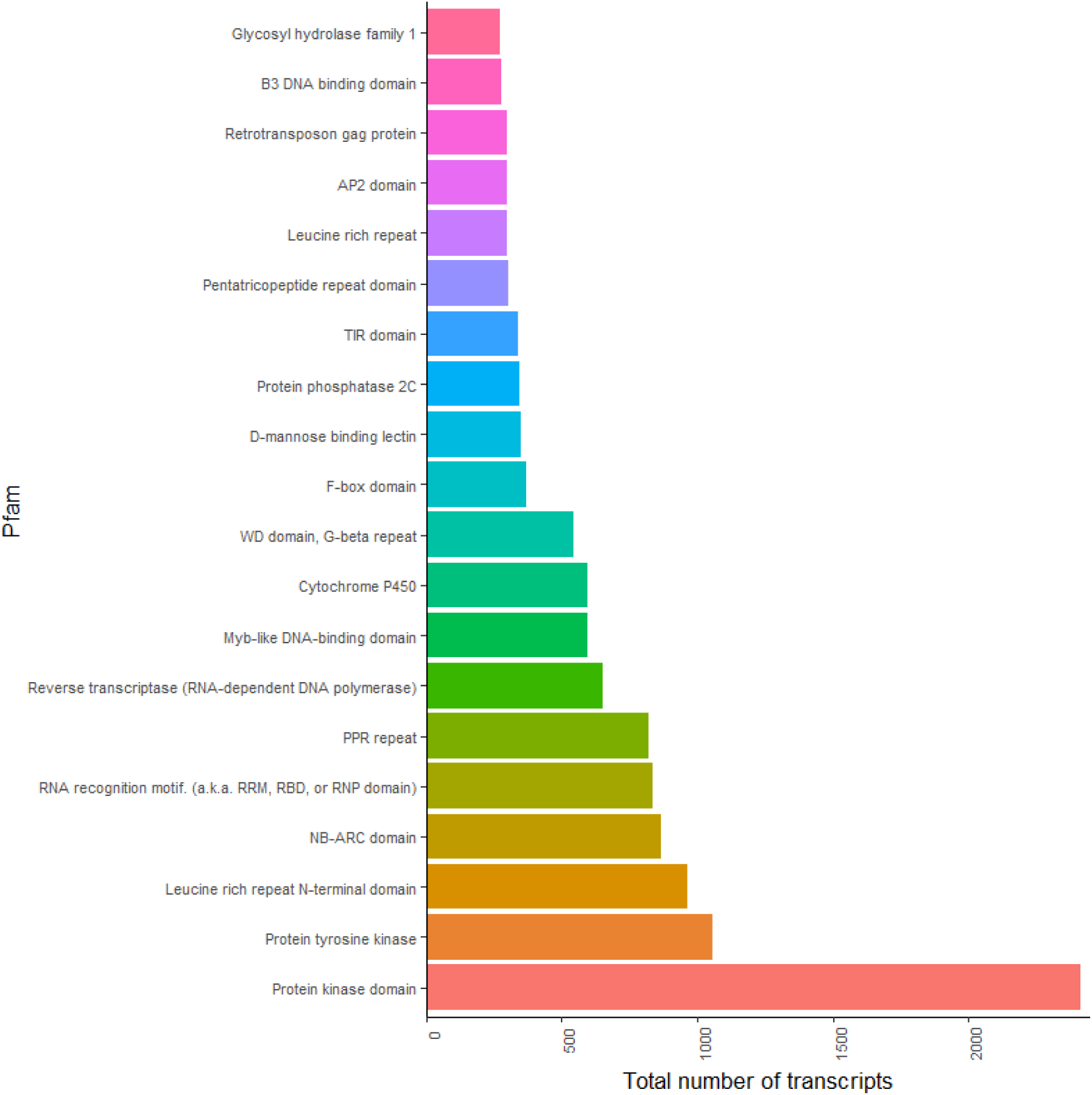
Top 20 PFAM domains in the comprehensive transcriptome.

The GO annotation retrieved a total of 58,500 terms for the Biological Process (BP) category, 61,467 terms for Molecular Function (MF) and 62,795 terms for Celular Component (CC). In addition, a total of 62,077 transcripts were annotated in the KEGG database.

### 3.3 Digital Gene Expression Analysis

To investigate the chilling stress response strategy between the rubber tree genotypes, we performed a pairwise comparison for each time series between GT1 and RRIM600. Prior to exposing the plants to cold stress (0 h), 624 genes were up-regulated in RRIM600 relative to GT1 (RRIM600 0h x GT1 0h), while 732 genes were up-regulated in GT1 0h relative to RRIM600 0h. After 90 minutes of cold stress exposure, we identified 514 genes that were up-regulated in RRIM600 90 min compared to GT1 90 min and 854 up-regulated genes in GT1 90 min relative to RRIM600 90min. Moreover, a total of 569 genes and 1034 genes were up-regulated in RRIM600 after 12 h and 24 h of cold stress exposure, respectively. Nevertheless, the GT1 genotype exhibited 610 and 875 up-regulated genes relative to RRIM600 after 12 h and 24 h of cold treatment, respectively (Figures 2 and 3).

**Figure 2.**
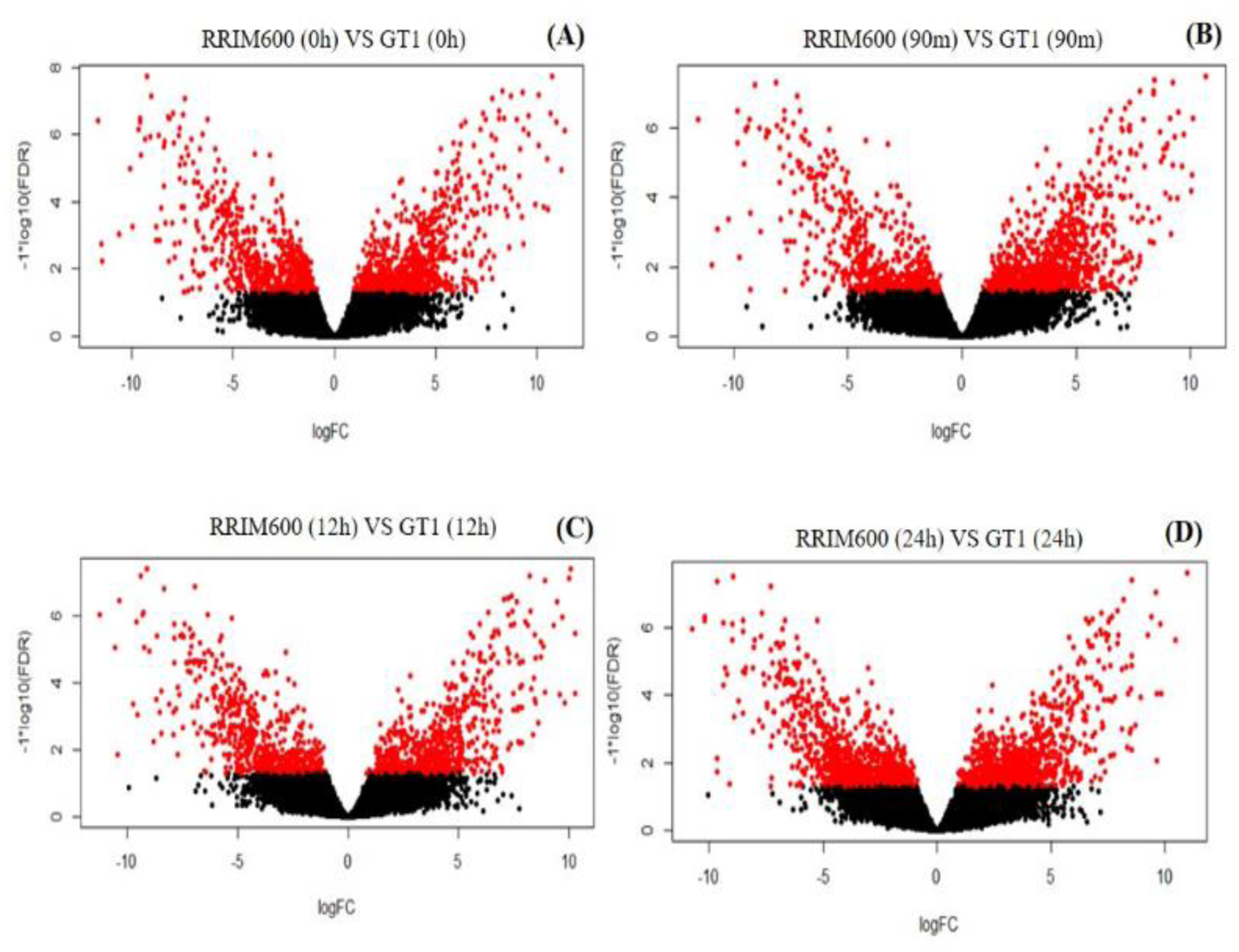
Volcano Plot of pair-wise comparison between RRIM600 and GT1 for each time-series of the cold treatment.

**Figure 3.**
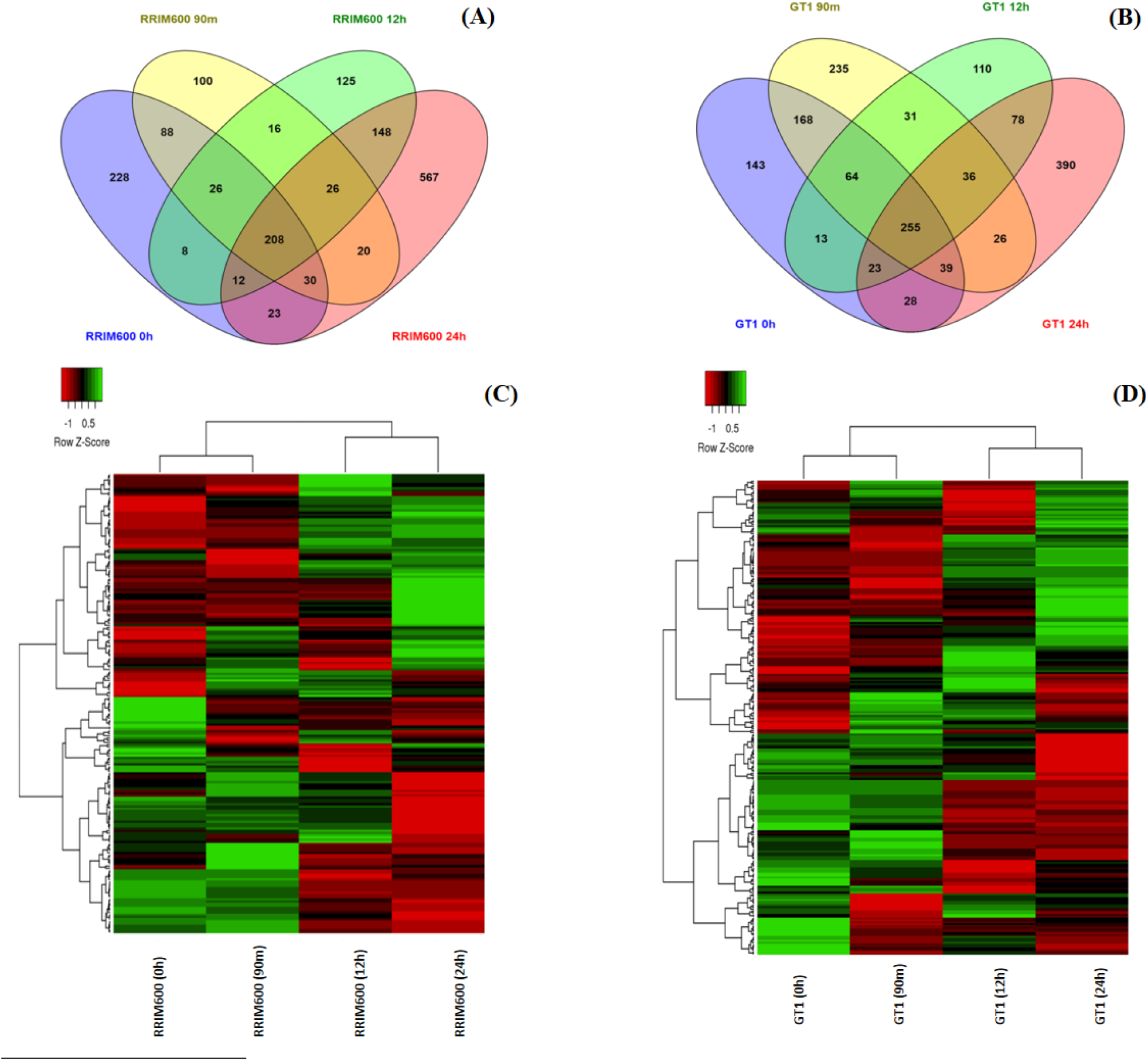
Expression profile for RRIM600 and GT1 genotypes. (A) Venn diagram representing the up-regulated genes identified in RRIM600 throughout the chilling treatment. (**B**) Venn diagram representing the up-regulated genes identified in GT1 throughout the chilling treatment. (**C**) Hierarchical clustering of DEGs for the 208 common overexpressed genes in RRIM600. (**D**) Hierarchical clustering of DEG of the 255 common overexpressed genes in GT1

Since the DEG analysis was performed by comparing the RRIM600 and GT1 genotypes for each time point of cold treatment, we compared the up-regulated genes previously identified for each genotype across all time points in order to identify genes that were commonly and exclusively up-regulated in the relative comparison between each treatment time. For the RRIM600 genotype, we detected a total of 229 genes that were exclusively up-regulated at 0 h (Figure 3A). After 90 minutes, 12 h and 24 h, we identified 100, 125 and 567 exclusively up-regulated genes, respectively. Moreover, a total of 208 RRIM600 genes were commonly up-regulated within the entire series (Figure 3A and C).

Whereas, there were a total of 143 up-regulated genes identified in GT1 relative to RRIM600 that were exclusively up-regulated at time 0 h and 235, 110 and 390 genes that were exclusively up-regulated after 90 minutes, 12 h and 24 h, respectively (Figure 3B). Furthermore, a total of 255 genes were commonly up-regulated across the time series (Figure 3B and D).

One of the genes up-regulated in RRIM600 relative to GT1 after 90minutes of cold stress was identified as a putative stelar K^+^ ‘outward-rectifying’ channel (SKOR) based upon a high BLAST similarity. Furthermore, two SKOR genes were up-regulated after 12 and 24 h of cold stress, one of which was among the ten most highly expressed genes after 12 h of treatment (Table 3, Supplementary Table 2). Reactive oxygen species (ROS) were recently shown to be capable of activating SKOR genes, thereby catalyzing K+ efflux from plant cells. In moderate stress conditions, K+ efflux could play the role of a ‘metabolic switch’ in anabolic reactions, stimulating catabolic processes and saving energy for adaption and repair (Demidchik et al., 2014).

**Table 3.**
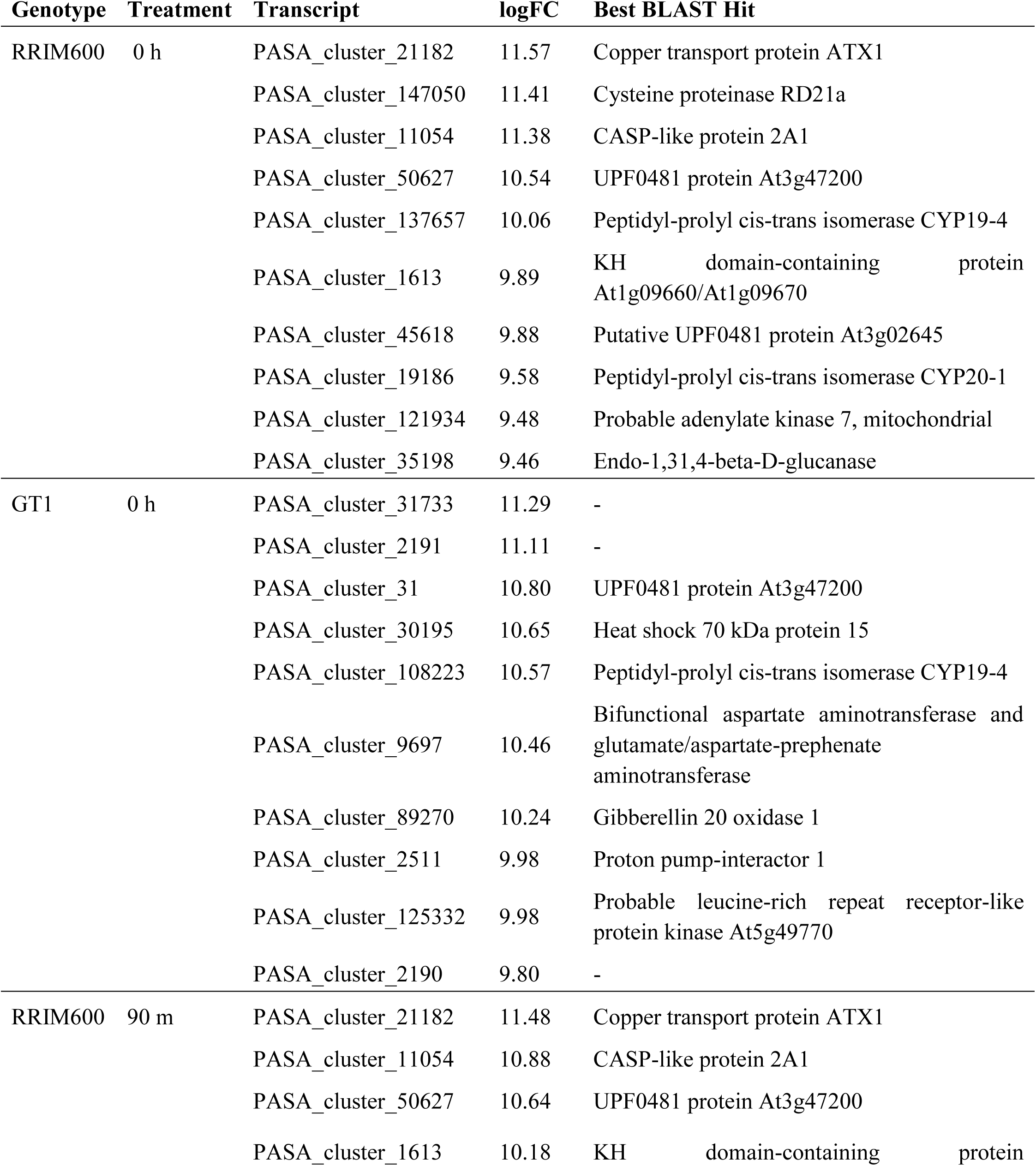
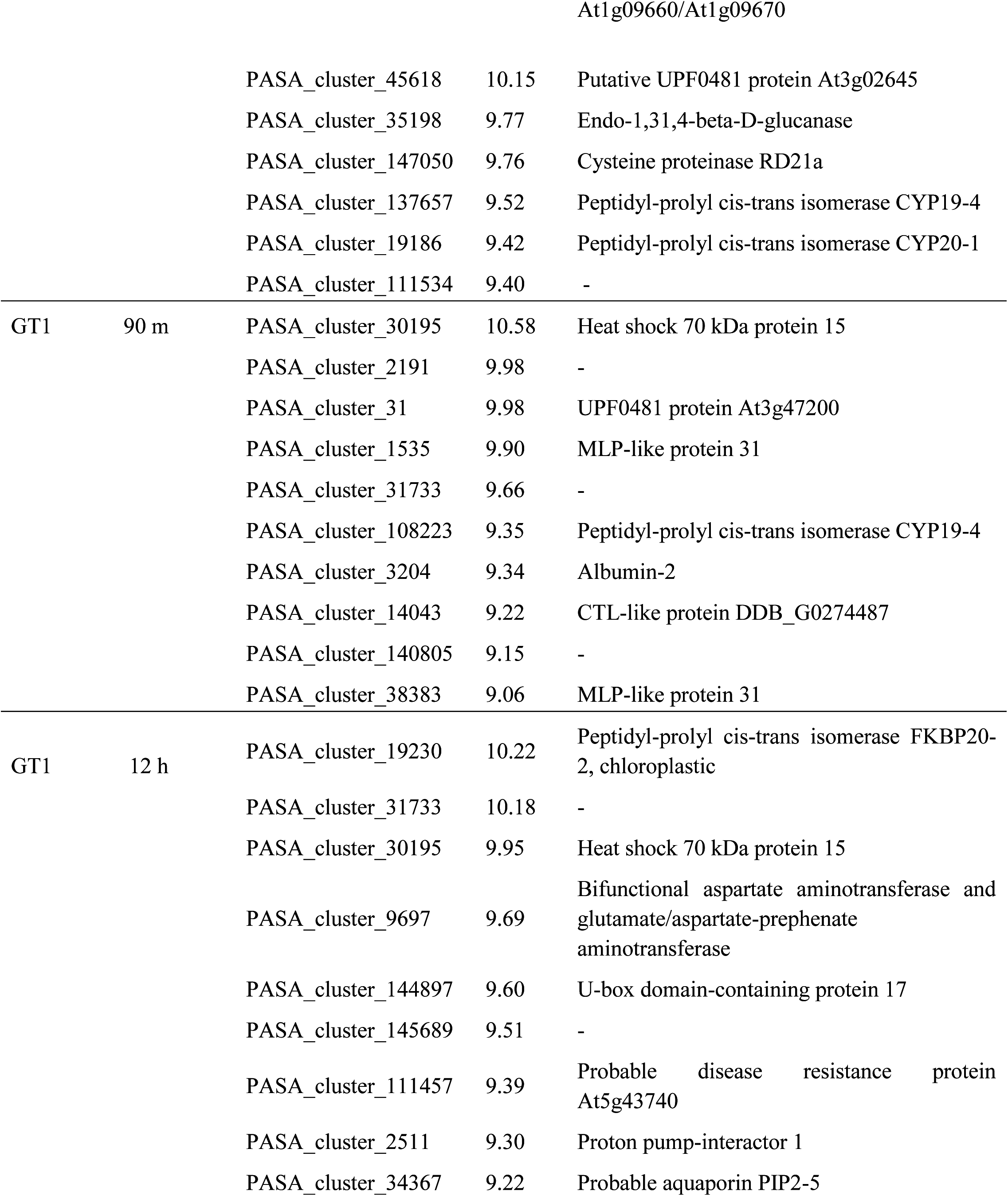
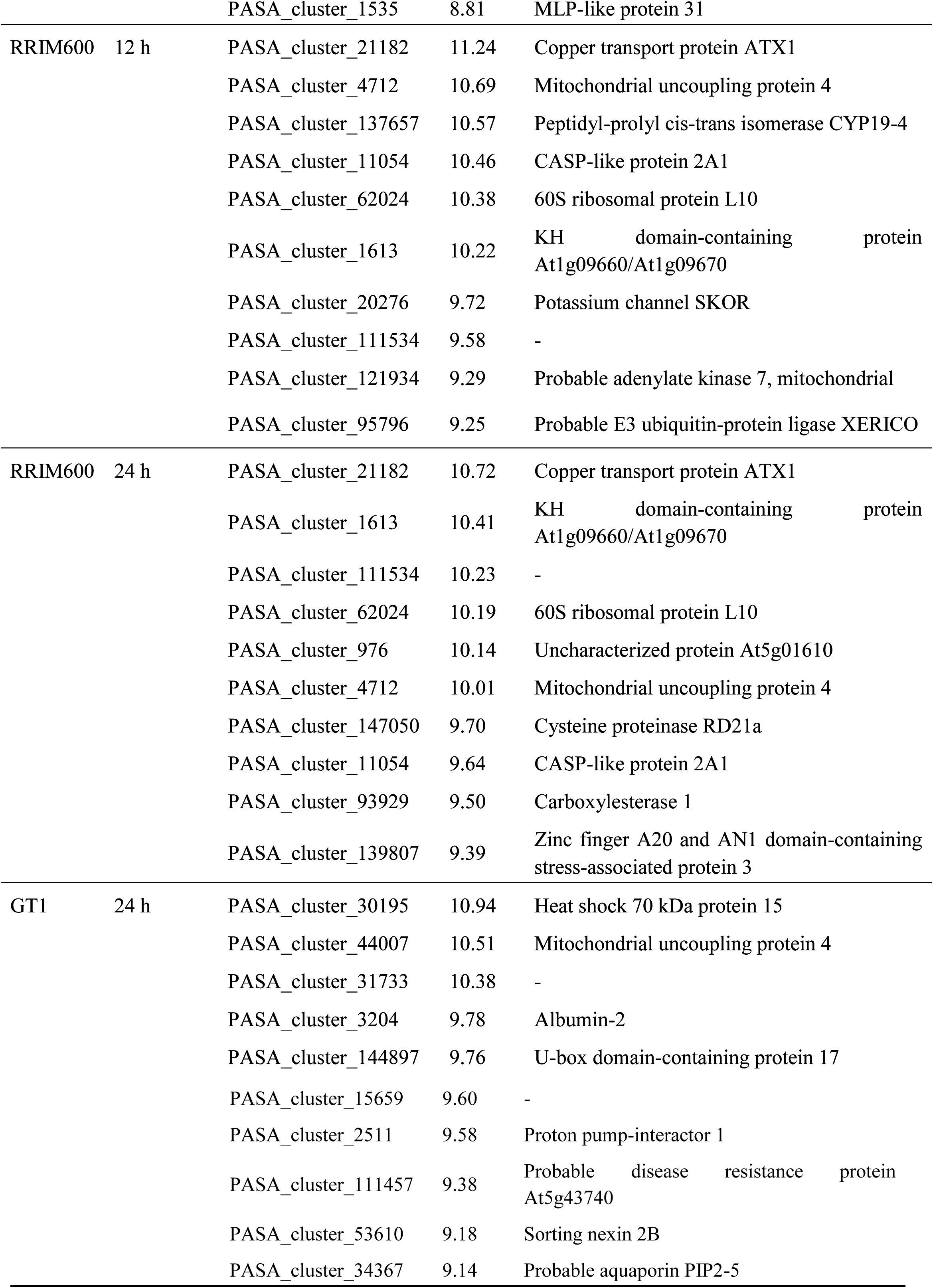
Top 10 overexpressed genes for each genotype in each time point.

| **Genotype** | **Treatment** | **Transcript** | **logFC** | **Best BLAST Hit** |
| --- | --- | --- | --- | --- |
| RRIM600 | 0 h | PASA_cluster_21182 | 11.57 | Copper transport protein ATX1 |
|  |  | PASA_cluster_147050 | 11.41 | Cysteine proteinase RD21a |
|  |  | PASA_cluster_11054 | 11.38 | CASP-like protein 2A1 |
|  |  | PASA_cluster_50627 | 10.54 | UPF0481 protein At3g47200 |
|  |  | PASA_cluster_137657 | 10.06 | Peptidyl-prolyl cis-trans isomerase CYP19-4 |
|  |  | PASA_cluster_1613 | 9.89 | KH domain-containing protein At1g09660/At1g09670 |
|  |  | PASA_cluster_45618 | 9.88 | Putative UPF0481 protein At3g02645 |
|  |  | PASA_cluster_19186 | 9.58 | Peptidyl-prolyl cis-trans isomerase CYP20-1 |
|  |  | PASA_cluster_121934 | 9.48 | Probable adenylate kinase 7, mitochondrial |
|  |  | PASA_cluster_35198 | 9.46 | Endo-1,31,4-beta-D-glucanase |
| GT1 | 0 h | PASA_cluster_31733 | 11.29 | - |
|  |  | PASA_cluster_2191 | 11.11 | - |
|  |  | PASA_cluster_31 | 10.80 | UPF0481 protein At3g47200 |
|  |  | PASA_cluster_30195 | 10.65 | Heat shock 70 kDa protein 15 |
|  |  | PASA_cluster_108223 | 10.57 | Peptidyl-prolyl cis-trans isomerase CYP19-4 |
|  |  | PASA_cluster_9697 | 10.46 | Bifunctional aspartate aminotransferase and glutamate/aspartate-prephenate aminotransferase |
|  |  | PASA_cluster_89270 | 10.24 | Gibberellin 20 oxidase 1 |
|  |  | PASA_cluster_2511 | 9.98 | Proton pump-interactor 1 |
|  |  | PASA_cluster_125332 | 9.98 | Probable leucine-rich repeat receptor-like protein kinase At5g49770 |
|  |  | PASA_cluster_2190 | 9.80 | - |
| RRIM600 | 90 m | PASA_cluster_21182 | 11.48 | Copper transport protein ATX1 |
|  |  | PASA_cluster_11054 | 10.88 | CASP-like protein 2A1 |
|  |  | PASA_cluster_50627 | 10.64 | UPF0481 protein At3g47200 |
|  |  | PASA_cluster_1613 | 10.18 | KH domain-containing protein |
| At1g09660/At1g09670 |  |  |  |  |
|  |  | PASA_cluster_45618 | 10.15 | Putative UPF0481 protein At3g02645 |
|  |  | PASA_cluster_35198 | 9.77 | Endo-1,31,4-beta-D-glucanase |
|  |  | PASA_cluster_147050 | 9.76 | Cysteine proteinase RD21a |
|  |  | PASA_cluster_137657 | 9.52 | Peptidyl-prolyl cis-trans isomerase CYP19-4 |
|  |  | PASA_cluster_19186 | 9.42 | Peptidyl-prolyl cis-trans isomerase CYP20-1 |
|  |  | PASA_cluster_111534 | 9.40 | - |
| GT1 | 90 m | PASA_cluster_30195 | 10.58 | Heat shock 70 kDa protein 15 |
|  |  | PASA_cluster_2191 | 9.98 | - |
|  |  | PASA_cluster_31 | 9.98 | UPF0481 protein At3g47200 |
|  |  | PASA_cluster_1535 | 9.90 | MLP-like protein 31 |
|  |  | PASA_cluster_31733 | 9.66 | - |
|  |  | PASA_cluster_108223 | 9.35 | Peptidyl-prolyl cis-trans isomerase CYP19-4 |
|  |  | PASA_cluster_3204 | 9.34 | Albumin-2 |
|  |  | PASA_cluster_14043 | 9.22 | CTL-like protein DDB_G0274487 |
|  |  | PASA_cluster_140805 | 9.15 | - |
|  |  | PASA_cluster_38383 | 9.06 | MLP-like protein 31 |
| GT1 | 12 h | PASA_cluster_19230 | 10.22 | Peptidyl-prolyl cis-trans isomerase FKBP20-2, chloroplastic |
|  |  | PASA_cluster_31733 | 10.18 | - |
|  |  | PASA_cluster_30195 | 9.95 | Heat shock 70 kDa protein 15 |
|  |  | PASA_cluster_9697 | 9.69 | Bifunctional aspartate aminotransferase and glutamate/aspartate-prephenate aminotransferase |
|  |  | PASA_cluster_144897 | 9.60 | U-box domain-containing protein 17 |
|  |  | PASA_cluster_145689 | 9.51 | - |
|  |  | PASA_cluster_111457 | 9.39 | Probable disease resistance protein At5g43740 |
|  |  | PASA_cluster_2511 | 9.30 | Proton pump-interactor 1 |
|  |  | PASA_cluster_34367 | 9.22 | Probable aquaporin PIP2-5 |
|  |  | PASA_cluster_1535 | 8.81 | MLP-like protein 31 |
| RRIM600 | 12 h | PASA_cluster_21182 | 11.24 | Copper transport protein ATX1 |
|  |  | PASA_cluster_4712 | 10.69 | Mitochondrial uncoupling protein 4 |
|  |  | PASA_cluster_137657 | 10.57 | Peptidyl-prolyl cis-trans isomerase CYP19-4 |
|  |  | PASA_cluster_11054 | 10.46 | CASP-like protein 2A1 |
|  |  | PASA_cluster_62024 | 10.38 | 60S ribosomal protein L10 |
|  |  | PASA_cluster_1613 | 10.22 | KH domain-containing protein<br>At1g09660/At1g09670 |
|  |  | PASA_cluster_20276 | 9.72 | Potassium channel SKOR |
|  |  | PASA_cluster_111534 | 9.58 | - |
|  |  | PASA_cluster_121934 | 9.29 | Probable adenylate kinase 7, mitochondrial |
|  |  | PASA_cluster_95796 | 9.25 | Probable E3 ubiquitin-protein ligase XERICO |
| RRIM600 | 24 h | PASA_cluster_21182 | 10.72 | Copper transport protein ATX1 |
|  |  | PASA_cluster_1613 | 10.41 | KH domain-containing protein<br>At1g09660/At1g09670 |
|  |  | PASA_cluster_111534 | 10.23 | - |
|  |  | PASA_cluster_62024 | 10.19 | 60S ribosomal protein L10 |
|  |  | PASA_cluster_976 | 10.14 | Uncharacterized protein At5g01610 |
|  |  | PASA_cluster_4712 | 10.01 | Mitochondrial uncoupling protein 4 |
|  |  | PASA_cluster_147050 | 9.70 | Cysteine proteinase RD21a |
|  |  | PASA_cluster_11054 | 9.64 | CASP-like protein 2A1 |
|  |  | PASA_cluster_93929 | 9.50 | Carboxylesterase 1 |
|  |  | PASA_cluster_139807 | 9.39 | Zinc finger A20 and AN1 domain-containing<br>stress-associated protein 3 |
| GT1 | 24 h | PASA_cluster_30195 | 10.94 | Heat shock 70 kDa protein 15 |
|  |  | PASA_cluster_44007 | 10.51 | Mitochondrial uncoupling protein 4 |
|  |  | PASA_cluster_31733 | 10.38 | - |
|  |  | PASA_cluster_3204 | 9.78 | Albumin-2 |
|  |  | PASA_cluster_144897 | 9.76 | U-box domain-containing protein 17 |

|  |  |  |
| --- | --- | --- |
| PASA_cluster_15659 | 9.60 | - |
| PASA_cluster_2511 | 9.58 | Proton pump-interactor 1 |
| PASA_cluster_111457 | 9.38 | Probable disease resistance protein<br>At5g43740 |
| PASA_cluster_53610 | 9.18 | Sorting nexin 2B |
| PASA_cluster_34367 | 9.14 | Probable aquaporin PIP2-5 |

Aquaporins belong to a conserved group of transmembrane proteins involved in water transport across membranes (King et al., 2004). GT1 contained an up-regulated gene with high similarity to the aquaporin subfamily plasma membrane intrinsic protein 2;5 (PIP2;5). The PIP2;5 gene was up-regulated across all time points in GT1 and was among the top ten up-regulated genes after 12 h and 24 h of cold treatment (Table 3, Supplementary Table 2).

A putative soybean gene regulated by cold 2 (SRC2) was up-regulated in RRIM660 across all time points. SRC2 is believed to play a role in cold stress responses in Arabidopsis (Kawarazaki et al., 2013) and soybean (Takahashi and Shimosaka, 1997). Notably, the log-fold-change in RRIM600 24h relative to GT1 24h was greater than the log-fold-change at earlier periods of cold treatment (Supplementary Table 2).

We also observed up-regulated genes in GT1 24h relative to RRIM600 24h, which were related to cell growth, such as the gene with high similarity to the LONGIFOLIA 1 (LNG1) protein, which in association with LONGIFOLIA 2 (LNG2) regulates leaf morphology by promoting longitudinal polar cell elongation (Lee et al., 2006). At the same time point in GT1 (24 h), we identified one up-regulated gene displaying high similarity to the RETICULATA-RELATED 6 (RER 6) protein. The RER6 gene may play a role in leaf development (Lee et al., 2006). Within the set of genes up-regulated in GT1 24h we identified a putative purine-uracil permease NCS1, which contributes to uracil import into plastids and is essential for plant growth and development (Mourad et al., 2012); and the receptor-like kinase TMK3, which is involved in auxin signal transduction and cell expansion and proliferation regulation (Dai et al., 2013) (Supplementary Table 2).

### 3.4 Protein Domain Homology among DEGs

Prior to cold stress, we detected four genes up-regulated in RRIM600 0h relative to GT1 0h with the Apetala 2 (AP2) domain. The AP2 genes show high similarity to ERF119, which may be involved in the regulation of gene expression by stress factors and by components of stress signal transduction pathways. The IQ calmodulin-binding motif domain was detected in six up-regulated genes in GT1 0h relative to RRIM600 0h. At 90 minutes, the number of up-regulated genes in GT1 containing this domain increased to eight (Table 4). The IQ calmodulin-binding is a major calcium (Ca^2+^) sensor and orchestrator of regulatory events through its interaction with a diverse group of cellular proteins (Rhoads and Friedberg, 1997) (Supplementary Table 2).

**Table 4.** The 10 most representative Pfam protein domain annotations for DEGs.

| Genotype | Treatment | Number of Genes | Pfam Domain |
| --- | --- | --- | --- |
| RRIM600 | 0 h | 13 | Protein tyrosine kinase |
|  |  | 11 | NB-ARC domain |
|  |  | 8 | Protein kinase domain |
|  |  | 7 | Plant protein of unknown function |
|  |  | 6 | Cytochrome P450 |
|  |  | 6 | Leucine rich repeat N-terminal domain |
|  |  | 6 | PPR repeat |
|  |  | 6 | RNA recognition motif. (a.k.a. RRM, RBD, o |
|  |  | 5 | Ribosomal protein L7Ae/L30e/S12e/Gadd45 fa |
|  |  | 4 | ADP-ribosylation factor family |
|  |  | 4 | AP2 domain |
|  |  | 4 | D-mannose binding lectin |
|  |  | 4 | KH domain |
|  |  | 4 | Leucine Rich repeat |
|  |  | 4 | Ribosomal protein L11, N-terminal domain |
|  |  | 4 | Ribosomal protein S8 |
| GT1 | 0 h | 16 | Leucine rich repeat N-terminal domain |
|  |  | 12 | Cytochrome P450 |
|  |  | 12 | Protein kinase domain |
|  |  | 11 | NB-ARC domain |
|  |  | 10 | PMR5 N-terminal domain |
|  |  | 8 | Ankyrin repeats (3 copies) |
|  |  | 8 | Glycosyl transferase family 8 |
|  |  | 7 | Microtubule binding |
|  |  | 6 | IQ calmodulin-binding motif |
|  |  | 6 | Legume lectin domain |
|  |  | 6 | Multicopper oxidase |
|  |  | 6 | Zinc-binding RING-finger |
|  |  | 5 | Myb-like DNA-binding domain |
|  |  | 5 | Peptidase inhibitor I9 |
|  |  | 5 | Short chain dehydrogenase |
|  |  | 5 | TIR domain |
| RRIM600 | 90 m | 13 | NB-ARC domain |
|  |  | 13 | Protein tyrosine kinase |
|  |  | 9 | Leucine rich repeat N-terminal domain |
|  |  | 8 | Protein kinase domain |
|  |  | 6 | D-mannose binding lectin |
|  |  | 5 | Leucine Rich repeat |
|  |  | 5 | Plant protein of unknown function |
|  |  | 4 | Cytochrome P450 |
|  |  | 4 | UDP-glucuronosyl and UDP-glucosyl transferase |
|  |  | 3 | Ammonium Transporter Family |
|  |  | 3 | CCAAT-binding transcription factor (CBF-B/NF-YA) subunit B |
|  |  | 3 | Leucine rich repeat |
|  |  | 3 | TIR domain |
|  |  | 3 | Wall-associated receptor kinase C-terminal |
|  |  | 3 | Wall-associated receptor kinase galacturonan-binding |
| GT1 | 90 m | 25 | Protein kinase domain |
|  |  | 22 | Leucine rich repeat N-terminal domain |
|  |  | 15 | NB-ARC domain |
|  |  | 13 | PMR5 N-terminal domain |
|  |  | 12 | Cytochrome P450 |
|  |  | 11 | Zinc-binding RING-finger |
|  |  | 9 | Ankyrin repeats (3 copies) |
|  |  | 9 | Glycosyl transferase family 8 |
|  |  | 8 | IQ calmodulin-binding motif |
|  |  | 7 | Microtubule binding |
|  |  | 7 | Short chain dehydrogenase |
| RRIM600 | 12 h | 17 | NB-ARC domain |
|  |  | 10 | Protein tyrosine kinase |
|  |  | 9 | Leucine rich repeat N-terminal domain |
|  |  | 7 | Protein kinase domain |
|  |  | 5 | F-box domain |
|  |  | 5 | Plant protein of unknown function |
|  |  | 5 | TIR domain |
|  |  | 5 | UDP-glucuronosyl and UDP-glucosyl transfer |
|  |  | 5 | VQ motif |
|  |  | 4 | Cytochrome P450 |
|  |  | 4 | D-mannose binding lectin |
|  |  | 4 | Leucine Rich repeat |
|  |  | 4 | PPR repeat |
| GT1 | 12 h | 25 | Protein kinase domain |
|  |  | 22 | Leucine rich repeat N-terminal domain |
|  |  | 15 | NB-ARC domain |
|  |  | 13 | PMR5 N-terminal domain |
|  |  | 12 | Cytochrome P450 |
|  |  | 11 | Zinc-binding RING-finger |
|  |  | 9 | Ankyrin repeats (3 copies) |
|  |  | 9 | Glycosyl transferase family 8 |
|  |  | 8 | IQ calmodulin-binding motif |
|  |  | 7 | Microtubule binding |
|  |  | 7 | Short chain dehydrogenase |
| RRIM600 | 24 h | 17 | NB-ARC domain |
|  |  | 15 | UDP-glucuronosyl and UDP-glucosyl transferase |
|  |  | 14 | Cytochrome P450 |
|  |  | 14 | PPR repeat |
|  |  | 12 | Protein kinase domain |
|  |  | 11 | Leucine rich repeat N-terminal domain |
|  |  | 11 | Protein tyrosine kinase |
|  |  | 9 | D-mannose binding lectin |
|  |  | 9 | Short chain dehydrogenase |
|  |  | 8 | AP2 domain |
| GT1 | 24 h | 23 | Leucine rich repeat N-terminal domain |
|  |  | 21 | Protein kinase domain |
|  |  | 20 | NB-ARC domain |
|  |  | 11 | TIR domain |
|  |  | 9 | Protein tyrosine kinase |
|  |  | 8 | Ankyrin repeats (3 copies) |
|  |  | 8 | Cytochrome P450 |
|  |  | 7 | Microtubule binding |
|  |  | 7 | PMR5 N-terminal domain |
|  |  | 6 | Carbohydrate-binding protein of the ER |
|  |  | 6 | Salt stress response/antifungal |

We also identified five up-regulated genes in RRIM600 12h compared to GT1 12h containing the VQ motif (Table 4). This domain is a plant-specific domain characteristic of a class of proteins that regulates diverse developmental processes, including responses to biotic and abiotic stresses, seed development, and photomorphogenesis (Jing and Lin, 2015).

After 24 h of cold treatment, the most abundant domain among the up-regulated genes in RRIM600 24h was the NB-ARC domain, which is a signaling motif that is shared by plant resistance products and is a regulator of cell death in animals (van der Biezen and Jones, 1998). The next two most common domains were UDP-glucuronosyl/UDP-glucosyl transferase, which catalyzes the transfer of sugars to a wide range of acceptor molecules and regulates their activities (Ross et al., 2001) followed by cytochrome P450 (CYP450). The CYP450 gene catalyzes diverse reactions leading to the precursors of structural macromolecules such as lignin and cutin, or is involved in the biosynthesis or catabolism of hormone and signaling molecules, such as antioxidants and defense compounds (Werck-Reichhart et al., 2002) (Table 4). Furthermore, eight up-regulated genes containing the AP2 domain were identified in RRIM600 24h relative to GT24h.

LRR N-terminal domains were the most abundant type of domain in GT1 24h up-regulated genes, followed by protein kinase domains and NB-ARC domains. The LRR N-terminal domain is involved in a number of biological processes, including cell adhesion, signal transduction, immune responses, apoptosis and disease resistance (Rothberg et al., 1990). Moreover, six up-regulated genes with a salt stress response/antifungal domain were identified in GT1 24h (Table 4).

Interestingly, we identified two up-regulated genes containing cold shock domain-containing protein 3 (CSP3) domain that were up-regulated in RRIM600 0 h. This protein domain shares a cold shock domain with bacterial CSPs and is involved in the acquisition of freezing tolerance in plants. In *Arabidopsis*, overexpression of these genes in transgenic plants confers enhanced freezing tolerance (Kim et al., 2009). We also identified one up-regulated gene with high similarity to the serine/threonine-protein kinase HT1 in RRIM600 0h (Supplementary Table 2). In *Arabidopsis*, the HT1 gene product has been reported to control stomatal movement in response to CO_2_ (Hashimoto et al., 2006).

In plants, the cold stress signal is transmitted to activate CBF-dependent (C-repeat/drought-responsive element binding factor-dependent) and CBF-independent transcriptional pathways, where the CBF-dependent pathway activates the CBF regulon (Chinnusamy et al., 2010). Some transcription factors, such as the ERF/AP2 factors, RAP2.1 and RAP2.6 and the C2H2-type zinc finger STZ/ZAT10, are cold response genes belonging to the CBF regulon (Fowler and Thomashow, 2002). In this study, we identified one up-regulated gene sharing high similarity with the transcription factor ZAT10 in RRIM600 after 90 min and 12 h of cold stress. Interestingly, GT1 contained a putative ZAT10 gene up-regulated only at 24h. (Supplementary Table 2).

The pentatricopeptide repeat (PPR) superfamily is one of the largest gene families in plants. For example, more than 400 members of this group have been identified in both rice and Arabidopsis (Yuan et al., 2009). Most of the PPR proteins are targeted to the chloroplast and mitochondria and are involved in many functions. They play important roles in response to developmental and environmental stresses. Additionally, a set of PPR genes has been reported to be involved in abiotic stress response regulation in Arabidopsis, through ROS homeostasis or ABA signaling (Zsigmond et al., 2008). In this study, we observed that the number of up-regulated PPR genes increased in RRIM600 and GT1 after 24 h of cold treatment. Prior to cold stress, RRIM600 showed six up-regulated genes, while GT1 had three up-regulated genes. However, after 24 h of chilling stress, RRIM600 revealed 14 up-regulated PPR genes, whereas GT1 presented seven (Supplementary Table 2).

### 3.4 DEG GO Enrichment

To identify enriched GO terms, we performed a GO enrichment analysis with the up-regulated genes identified in RRIM600 and GT1 for each time point (Supplementary Table 3). Among all terms identified for all up-regulated genes in each genotype, a total of 32 non-redundant Biological Process (BP) terms were identified across the time series in RRIM600. Furthermore, we detected a total of 20 and 31 distinctive terms in the Cellular Component (CC) and Molecular Function (MF) categories, respectively. However, GT1 presented a total of 102 non-redundant terms in the BP category. The CC and MF categories exhibited a total of 37 and 44 unique terms, respectively.

We observed enriched BP terms related to defense, such as the defense response (GO:0006952), cellular defense response (GO:0006968) and innate immune response (GO:0045087), in the up-regulated genes prior to cold stress (0h) in the RRIM600 genotype. Additionally, we observed a substantial increase in the number of sequences related to the defense response category. Before cold treatment, RRIM600 contained 71 up-regulated genes in this category. After 24 h, we observed a total of 97 genes associated with defense responses. The GT1 clone began to exhibit enriched stress responses categories related after 90 minutes of cold treatment, such as the defense response (GO:0006998) (Supplementary Table 3). Interestingly, we also observed that the GT1 gene set was enriched for the lignin biosynthetic process (GO:0009809) and lignin metabolic process (GO:0009808) at 90 m. Across all time points, GT1 showed enriched categories such as cell wall (GO:0042546), plant-type cell wall biogenesis (GO:0009832), plant-type cell wall organization or biogenesis (GO:0071669) and cell wall biogenesis (GO:0042546) (Supplementary Table 2). Under cold stress, the secondary cell wall may be reinforced by the incorporation of lignin, which strengthens the wall and impedes cell damage and water loss (Le Gall et al., 2015).

After 24 h at 10°C, the respiratory chain category was enriched in the up-regulated genes in RRIM600 24h (GO:0070469), whereas the up-regulated genes identified in GT1 24h exhibited enriched terms such as thylakoid part (GO:0044436), photosystem II (GO:0009523) and photosystem II oxygen evolving complex (GO:0009654) (Supplementary Table 2).

For the MF category, the number of up-regulated genes annotated with cellulose synthase activity (GO:0016759) increased from 10 to 14 for 0h and 90 min in GT1. Whereas, at 24 h of cold stress, the number of cellulose synthase genes decreased to 9. Interestingly, RRIM600 did not show any enriched categories related to cellulose synthase or categories that could be related to the lignification process during cold treatment.

Recent studies have indicated that the purine metabolite allantoin can be involved in the response to stress, for example, playing a role in ABA metabolism and jasmonic signaling and, hence, promotes cold tolerance in plants (Watanabe et al., 2014; Takagi et al., 2016). Examination of the cold stress response in Chinese yew showed that purine metabolism was up-regulated (Meng et al., 2017). In RRIM600, the categories purine ribonucleotide binding (GO:0032555), purine nucleotide binding (GO:0017076), purine nucleoside binding (GO:0001883) and purine ribonucleoside binding (GO:0032550) were enriched from 90 min to 24 h of chilling treatment. Furthermore, the number of up-regulated genes related to these GO categories increased during cold treatment (Supplementary Table 3).

### 3.5 Putative Molecular Marker Detection

#### Microsatellite Discovery

The comprehensive transcriptome was evaluated by searching for microsatellites. A total of 27,111 SSRs were found in 21,237 transcripts, and 4,570 transcripts contained more than 1 SSR per sequence. The SSR frequency in this transcriptome was 1 SSR per 7.2 Kb.

Among the total putative SSRs detected, 16,621 (61%) were classified as dinucleotides, followed by 9,336 (34%) tri-, 634 (2%) tetra-, 283 (1%) penta-and 237 (1%) hexanucleotides. Among the dinucleotide SSRs, the most abundant motif was AG/TC, at 12,075 (72.65%), followed by the AT/TA, AC/TG and GC/CG motifs, 2,939 (17.68%), 1,560 (9.38%) and 47 (0.2%), respectively (Table 5).

**Table 5.** Statistical summary of the results of searches for putative SSRs.

|  |  |
| --- | --- |
| Total SSRs | 27,111 |
| Dinucleotide | 16,621 |
| AG/TC | 12,075 |
| AT/TA | 2,939 |
| AC/TG | 1,560 |
| GC/CG | 47 |
| Trinucleotide | 9,336 |
| Tetranucleotide | 634 |
| Pentanucleotide | 283 |
| Hexanucleotide | 237 |
| SSR frequency | 1 per 7.2 Kb |

The up-regulated genes detected in the GT1 and RRIM600 genotypes were merged to identify putative SSRs. A total of 1,034 dinucleotide SSRs were identified, followed by 629 tri-,18 tetra-,19 penta-and 23 hexanucleotide SSRs (Table 5).

#### SNP Discovery

SNP calling was performed for each genotype using Freebayes. A total of 202,949 putative SNPs were detected in GT1. Transition (Ts) SNPs were more abundant compared with transversion (Tv) SNPs, resulting in a Ts/Tv ratio of 1.46. Among the Ts variations, A↔G was the most abundant, with 61,111 putative SNPs, while A↔T was the most abundant variation in the Tv SNPs, with 24,613 markers (Table 6). The SNP frequency for GT1 was 1 SNP per 967 bp.

**Table 6.**
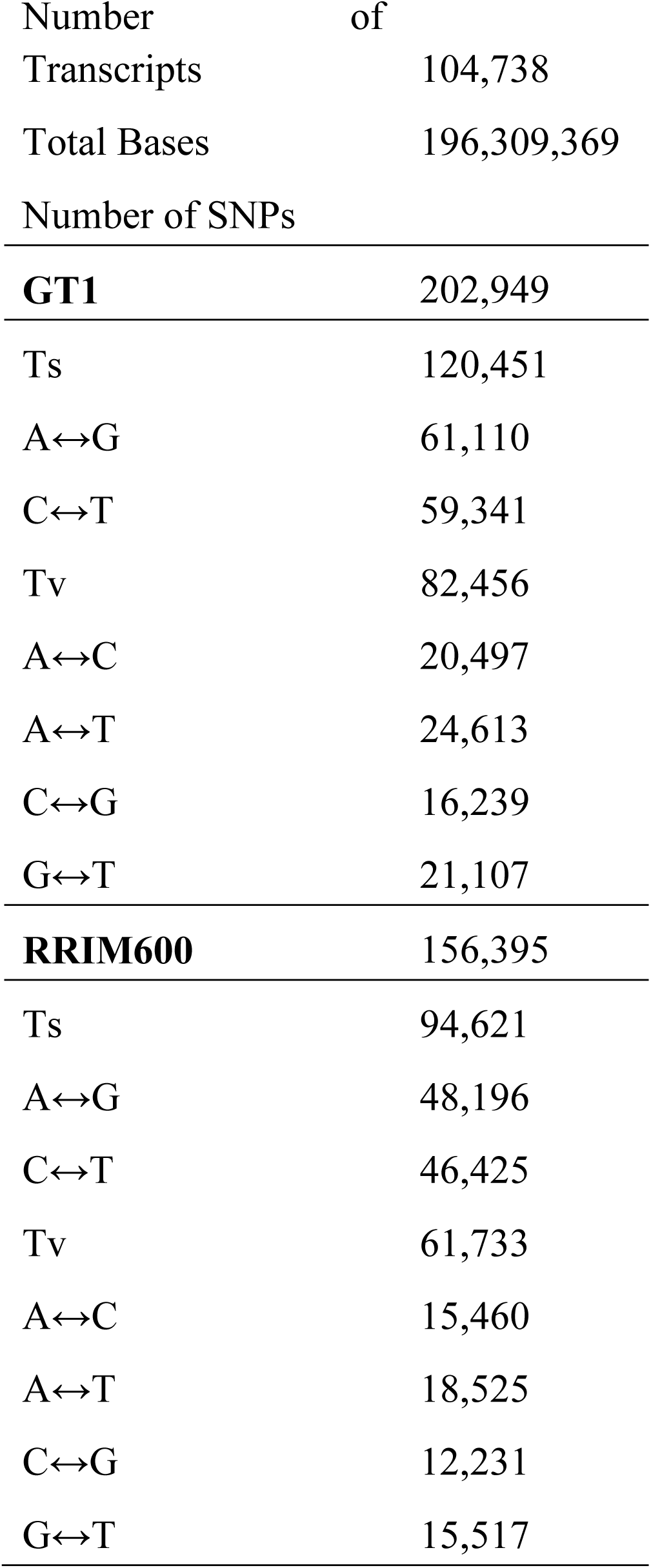
Statistical summary of the results of searches for putative SNPs.

For the RRIM600 genotype, a total of 156,354 putative SNPs were detected, and the Ts/Tv ratio was 1.53. As observed in GT1, A↔G was the most abundant variation, with 48,196 SNPs. The most frequent variation among the Tvs was A↔T, with 18,525 putative SNPs. The SNP frequency was 1 SNP per 1,255 kb (Table 6).

A total of 94,962 SNPs were common between GT1 and RRIM600, and the overall SNP frequency was 1 SNP per 742 bp. Among the DEGs, we identified 20,203 and 14,998 SNPs in GT1 and RRIM600, respectively. Among the SNPs identified in DEGs, 12,509 SNPs were exclusive to GT1 and 7,484 SNPs were exclusively to RRIM600.

### qRT-PCR Validation

To validate the DEG analysis, a total of 20 genes were selected, and primer pairs were designed to validate the analysis (Supplementary Table 1). All primer pairs were initially tested via PCR using genomic DNA as a template to verify the amplification product. From the 20 primer pairs, 14 were successfully amplified and used for qRT-PCR.

The qRT-PCR assays were conducted using RH2b and YSL8 as housekeeping genes. Among the 14 genes tested (Figure 4, Supplementary Table 4), 11 were differentially expressed between RRIM600 and GT1 and confirmed the *in silico* analysis. The gene encoding the DELLA protein GAI1 (PASA_cluster_35787) was detected in the *in silico* analysis as up-regulated in RRIM600 across all time point; however, this qPCR results revealed that gene was up-regulated at 0 h and 90 minutes of cold stress. For the HSP70 gene (PASA_cluster_30195), which was also identified as up-regulated across all time point in GT1 in the *in silico* analysis, the qPCR confirmed that the HSP70 gene was up-regulated for the 0 h, 90 min and 12 h time points. After 24 h, HSP70 levels were greater in GT1 than RRIM600, nevertheless this difference was not significant (Figure 4).

**Figure 4.**
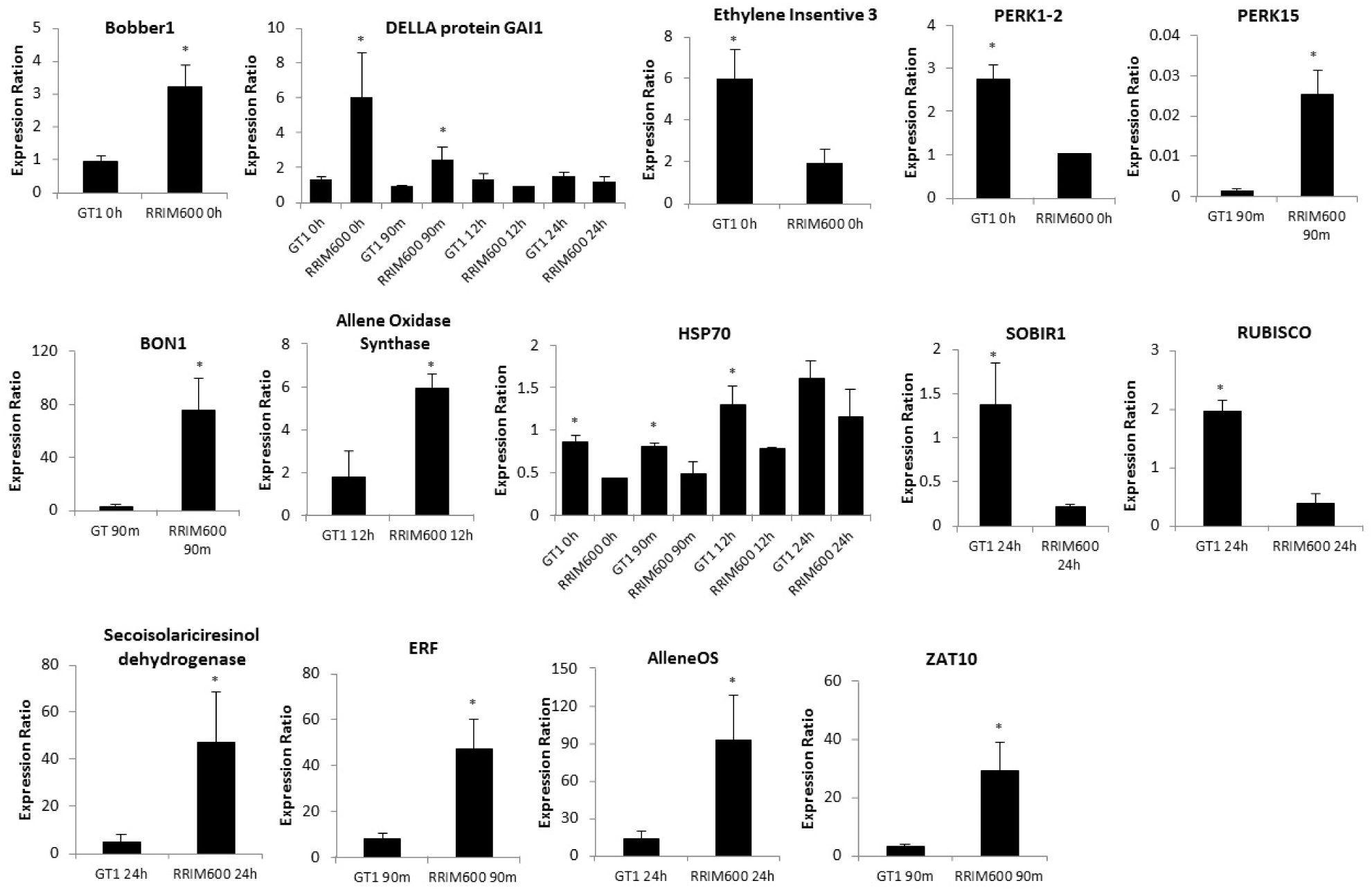
qPCR expression analysis of the 14 genes identified in the *in silico* DEG analysis. The expression values represent the mean (n=3 or n=2) ± SEM. The bars indicate the standard error of the mean values and significant differences (p < 0.05) are indicated with asterisks.

The only gene that is not in accordance with the *in silico* analysis was the protein ETHYLENE INSENSITIVE 3 (EIN3) (PASA_cluster_52015). The *in silico* analysis showed higher expression levels for EIN3 in RRIM600; however, qRT-PCR results showed that this gene is significantly up-regulated in GT1 (Figure 4).

#### Alternative Splicing Detection

AS is an important mechanism involved in gene regulation that may regulate many physiological processes in plants, including the response to abiotic stresses such as cold stress (Tack et al., 2014). In Arabidopsis, it has been estimated that 60% of genes are subject to AS (Filichkin et al., 2010). Furthermore, studies in soybean and maize predicted that 52% (Shen et al., 2014) and 40% (Thatcher et al., 2014) of genes are subject to AS events.

Due to the importance of AS, the reference transcriptome obtained in this study was also used to detect AS events. A minimum depth of 10 reads was used as the threshold to identify isoforms. A total of 20,279 AS events were identified, with intron retention (IR) representing the major AS event, accounting for a total of 9,226 events (45.5%), followed by exon skipping (ES), alternative acceptor (AltA) and alternative donor (AltD) events, at 4,806 (23.7%), 3,599 (17.7%) and 2,648 (13%) events, respectively (Figure 5).

**Figure 5.**
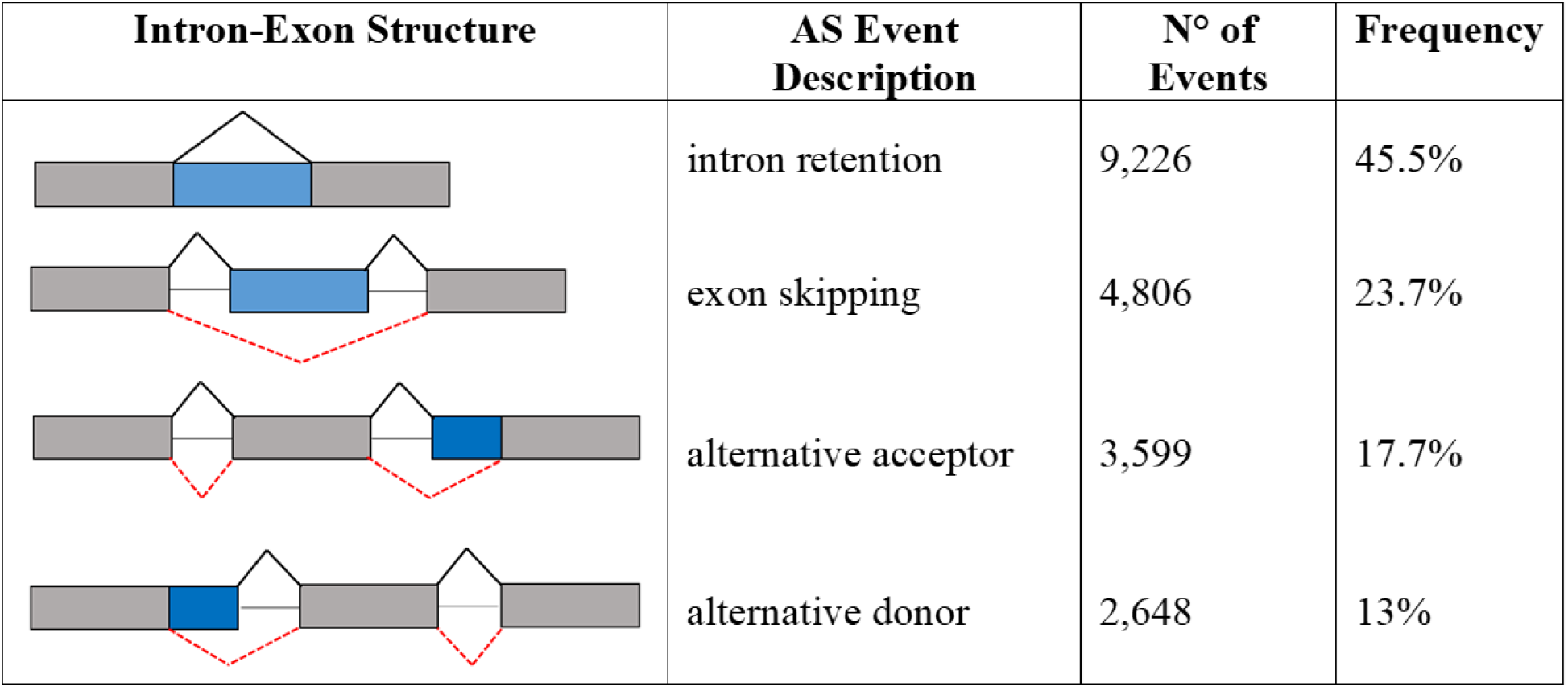
Summary of AS detection in the comprehensive transcriptome.

Although the ES type accounts for the majority of AS events in humans, it has been reported that IR events are the most abundant type in plants (Chamala et al., 2015).

## 4 Discussion

Abiotic stress is caused by environmental conditions such as cold and drought, consequently affecting optimum growth and yields. Crop production can be influenced by as much as 70% by environmental factors (Cramer et al., 2011). The inhibition of growth is one of the earliest responses to abiotic stress. The metabolism of lipids, sugar and photosynthesis is affected gradually as the stress becomes more severe. The plant response to abiotic stresses is complex and involves interactions and crosstalk with many molecular pathways (Cramer et al., 2011). Therefore, one of the stress tolerance mechanisms could be defined as the ability to detect stress factors and respond to them appropriately and efficiently (Sewelam et al., 2016).

### 4.1 Reactive Oxygen Species Scavenging

ROS are continuously produced at basal levels under favorable conditions. Organisms exhibit antioxidant mechanisms that scavenge ROS to maintain the appropriate balance (Foyer and Noctor, 2005). In recent years, it has been reported that ROS play an important signaling role in plants in response to biotic and abiotic stresses and in controlling processes such as growth (Das and Roychoudhury, 2014). However, different types of stress factors, such as drought, pathogen infection and extreme temperatures, disturb the balance between ROS generation and ROS scavenging, causing oxidative damage to membranes, proteins, RNA and DNA (Mittler, 2002).

The survival of plants therefore depends on many important factors, such as changes in growth conditions, the severity and duration of stress conditions and the capacity of the plants to quickly adapt to changing energy equations (Miller et al., 2010). Under stressful conditions, plant redox homeostasis is maintained by both antioxidant enzymes, such as pH-dependent peroxidases (POXs), superoxide dismutase (SOD), ascorbate peroxidase (APX), guaiacol peroxidase (GPX), glutathione-S-transferase (GST), and catalase (CAT), and non-enzymatic compounds, such as ascorbic acid (AA), reduced glutathione (GSH), α-tocopherol, carotenoids, phenolics, flavonoids, and proline (Gill and Tuteja, 2010; Miller et al., 2010).

Before exposing the plants to cold stress in the present study, both genotypes presented up-regulated genes with similarity to peroxidase. However, GT1 exhibited two up-regulated gene, while RRIM600 exhibited only one. After 90 minutes, only GT1 exhibited up-regulated peroxidase genes. At this time point, we detected a total of four up-regulated genes with the best BLAST hit to peroxidase. However, there were three up-regulated genes in RRIM600 with probable peroxidase activity after 24h of treatment (Figure 6, Supplementary Table 2).

**Figure 6.**
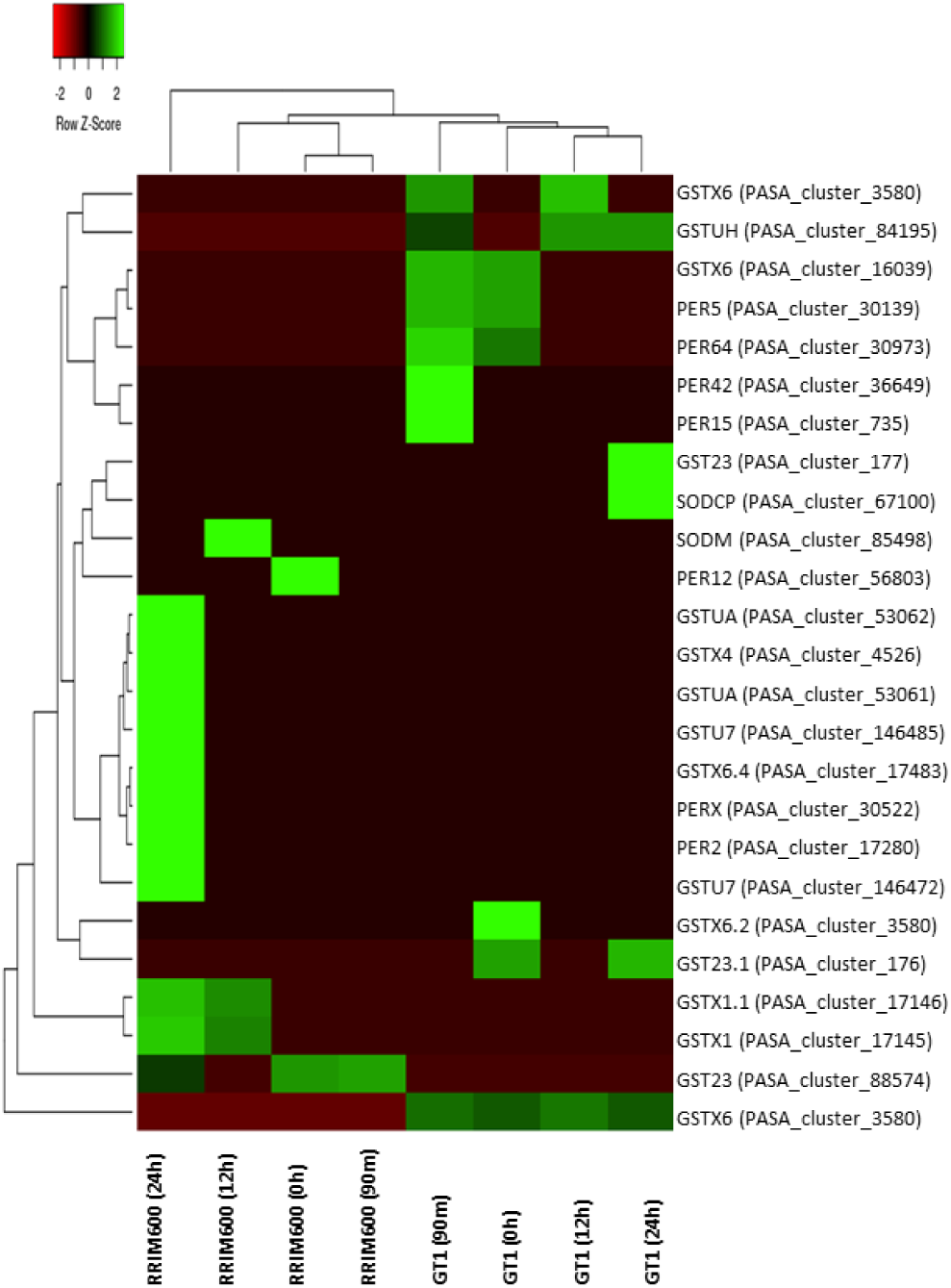
Heat Map for ROS scavenging indicating the up-regulated genes for each genotype in each time series.

SODs are the first line of defense of ROS (Alscher et al., 2002) and we identified two up-regulated putative SOD genes in both genotypes. The putative SOD gene that was up-regulated in RRIM600 was detected after 12 h of cold stress, while the SOD gene in GT1 was up-regulated after 24 h of treatment (Figure 6, Supplementary Table 2). Furthermore, putative GST genes were identified across the time series, where one GST gene was up-regulated in RRIM600 and three were up-regulated in GT1 before cold stress treatment (Figure 6). After 90 minutes, we detected five genes with high similarity to GST genes. Among these five genes, four were up-regulated in GT1 and one was up-regulated in RRIM600. At the subsequent time point, we observed an increase in up-regulated GST genes in RRIM600. After 24 hours, RRIM600 up-regulated nine putative GST genes, whereas GT1 had only four putative GST genes (Figure 6, Supplementary Table 3). The significant number of GST genes that were up-regulated in RRIM600 after 24h of cold treatment, indicates that GST genes may play an important role in ROS scavenging, thereby maintaining the integrity of cells.

Furthermore, we identified two up-regulated putative thioredoxin H1(TRXh1) genes in RRIM600 after 24 h of cold stress. In *Oryza sativa*, the TRXh1 gene is involved in stress responses where it regulates the balance of ROS in the rice apoplast. TRXh1 plays an important role in redox state regulation and stress responses (Zhang et al., 2011) (Supplementary Table 2).

### 4.2 Mitogen-Activated Protein Kinase (MAPK) Signaling Pathway

MAPK cascades are important signaling modules that transduce environmental signaling modules to transcriptional cascades (Zhao et al., 2017). The MEKK1-MKK2-MPK4/6 MAPK cascade participates in abiotic and biotic signaling to downstream ROS and plays a positive role in the cold stress response (Teige et al., 2004).

MEKK1/ANP1 has been reported to be activated under biotic and abiotic conditions such as cold, salt and drought (Teige et al., 2004), but it blocks the action of auxin, a plant mitogen and growth hormone (Kovtun et al., 2000). In Arabidopsis, constitutive expression of MEKK1 imitates the H_2_O_2_ effect and initiates a MAPK cascade that induces specific stress-responsive genes; however, this gene blocks the action of auxin. In tobacco, transgenic plants that constitutively overexpress MEKK1 show enhanced tolerance to multiple environmental stress conditions (Kovtun et al., 2000).

The MAP kinase kinase MKK2 phosphorylates the MPK4 and MPK6 MAP kinases in response to cold and salt stress (Teige et al., 2004). Additionally, plants overexpressing MKK2 showed constitutive MPK4 and MPK6 activity, which increases freezing and salt tolerance (Teige et al., 2004). Arabidopsis null mutants of mkk2 show compromised activation of MPK4 and MPK6 and are hypersensitive to cold and salt stress.

The KEGG annotation revealed up-regulated genes in the MAPK pathways for both genotypes across the time series. GT1 and RRIM600 exhibited up-regulated MKK2 and MPK4/6 putative genes before the initiation of the chilling treatment. However, pair-wise comparison between RRIM600 and GT1 revealed that RRIM600 exhibited one MEKK1/ANP1 (EC 2.7.11.25) gene that was exclusively uo-regulated after 90 minutes of cold stress.

Regarding MAPK signaling pathway, RRIM600 showed up-regulation of the basic endochitinase B (CHI-B) (EC 3.2.1.14) gene at 12 h and 24 h. This gene product is involved in the ethylene/jasmonic acid signaling pathway. Ethylene/jasmonic acid have also been shown to neutralize chilling stress, activate ROS scavenging enzymes, and regulate the C-repeat binding factor (CBF) pathway during cold stress (Sharma and Laxmi, 2016). In this study, the number of genes exhibiting high similarity to CHI-B increased from 1 to 4 after 24 h.

Interestingly, after 24 h of cold exposure, GT1 exhibited one up-regulated gene encoding catalase [CAT] isozyme 1, which catalyzes the decomposition of hydrogen peroxide (H_2_O_2_) and plays an important role in controlling the homeostasis of ROS (Du et al., 2008). At the same time point, GT1 also presented one up-regulated gene with cytoplasmic 5’-to-3’ exoribonuclease activity (EC 3.1.13.-) and one 1-aminocyclopropane-1-carboxylate synthase (ACS) gene (EC 4.4.1.14), which is the rate-limiting enzyme of ethylene biosynthesis. ACS is also induced by stress and a substrate of MPK6 (Liu & Zhang, 2004). These results are believed to indicate that the MAPK pathway is involved in ethylene signaling (Ma and Bohnert, 2007).

### 4.3 Signal Transduction

In plants, recognition of abiotic stress signals initiates specialized signaling pathways in which phosphatases and protein kinases are key components, such as CaM domain-containing protein kinases (CDPKs), calcineurin B-like proteins (CBLs), CBL-interacting protein kinases (CIPKs) and receptor-like kinases (RLKs), including LRR_RLK, MRLK and Lectin RLK (LecRLK) (Bose et al., 2011).

In this study, we detected a significant increase in up-regulated LecRLK genes in RRIM600 relative to GT1 after 24 h of cold treatment. Before initiating the cold treatment, the LecRLK gene was not up-regulated in RRIM600; however, after 24 h, we detected eight up-regulated genes. Interestingly, in GT1, the number of genes identified during cold treatment decreased from seven to one gene (Figure 7A, Supplementary Table 1). LecRLK genes were previously reported to enhance resistance to pathogen infection in tobacco (Wang et al., 2016) and Arabidopsis (Bouwmeester and Govers, 2009) and to a play role in abiotic stress signal transduction (Singh and Zimmerli, 2013).

**Figure 7.**
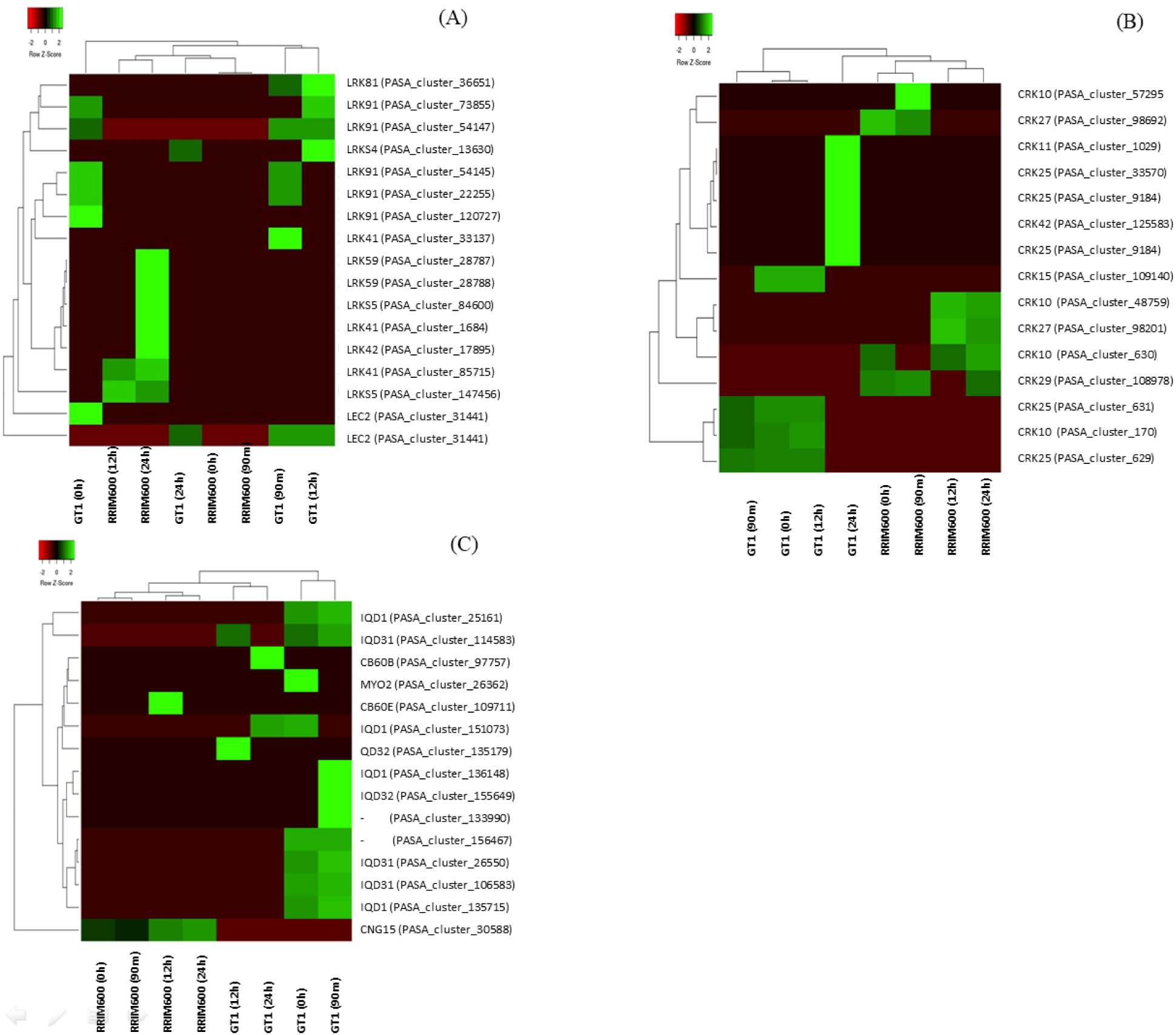
Heat Map with Legume Lectin-RLK (A) CRKs (B) and Calmodulin (C) up-regulated genes for each genotype in each time series.

Cold stress induces the elevation of cytosolic Ca^2+^ as an early response. This increase in cytosolic Ca^2+^ is suggested to be an important messenger for signal transduction and therefore cold acclimation. In alfalfa and Arabidopsis, a positive correlation between the cold-induced cytosolic Ca^2+^ increase and the accumulation of cold-induced transcripts has been observed (Monroy and Dhindsa, 1995; Henriksson and Trewavas, 2003). Ca^2+^ from the cytosol can be detected by Ca^2+^ sensors, such as calmodulin (CaM), CaM domain-containing protein kinases (CDPKs), calcineurin B-like proteins (CBLs) and CBL-interacting protein kinases (CIPKs) (Henriksson and Trewavas, 2003).

CaM genes were detected in both genotypes (Figure 7C). Relative to RRIM600, GT1 showed up-regulation of eight CaM genes prior to cold stress, while after 90 minutes, we identified a total of nine CaM genes (Figure 7C). We identified two up-regulated CaM putative genes in GT1 after 12 h and 24 h of cold stress. However, RRIM600 showed only one up-regulated CaM gene before cold treatment relative to GT1 (Figure 7C). This gene was up-regulated in RRIM600 during the entire treatment period. Interestingly, we also identified one up-regulated CIPK gene in GT1 at 90 minutes (Supplementary Table 2). Calcium/calmodulin-regulated receptor-like kinase 1 (CRLK1) is required for cold tolerance via the activation of MAP kinase activity (Yang et al., 2010). This gene activates MEKK1 in response to cold in a calcium-dependent manner (Furuya et al., 2013). The differential expression analysis between the two genotypes identified one CRLK1 in GT1 after 12 h of cold stress. Additionally, we identified one up-regulated CDPK2 gene in RRIM600 after 24 h of treatment.

Cysteine-rich receptor-like kinases (CRKs) can significantly affect plant development and stress responses (Burdiak et al., 2015). It has been suggested that CRK transcript levels are elevated in response to salicylic acid (SA), pathogens and drought. Additionally, CRKs are involved in mediating the effects of ROS (Bourdais et al., 2015). It was observed that the number of up-regulated CRK genes increased in both genotypes due to cold treatment. Before chilling stress, the comparison between GT1 and RRIM600 exhibited four and three up-regulated CRK genes, respectively. However, after 24 h cold treatment, GT1 and RRIM600 exhibited five and four up-regulated CRK genes, respectively. Interestingly, all of the up-regulated genes identified in GT1 after 24 h are different from the previous up-regulated CRK genes (Figure 7B, Supplementary Table 2).

Proline-rich extensin-like receptor protein kinase (PERK) is structurally organized into a proline-rich N-terminal domain, followed by a transmembrane segment and a C-terminal serine/threonine kinase domain (Florentino et al., 2006). The PERK genes are assigned to two major groups. One group is expressed exclusively in pollen, and the other is expressed throughout all tissues. PERKs may be involved in the early response to general perception and the response to wounding and/or pathogen stimuli and turgor pressure (Osakabe et al., 2014). GT1 0h and RRIM600 0h exhibited two and three up-regulated genes, respectively. During cold treatment, the number of up-regulated genes increased in both genotypes. After 24 h of chilling stress, the comparison between RRIM600 and GT1 revealed five and seven up-regulated genes in each genotype, respectively (Supplementary Table 2).

### 4.4 Photosynthesis Activity and Stomata Closure

Cold stress can cause an imbalance between light utilization and energy dissipation through metabolic activity. An excess of photosystem II (PSII) excitation pressure exists, which can be reversible through the dissipation of excess absorbed energy or irreversible inactivation of PSII and, consequently, inhibition of the photosynthetic capacity (Oquist and Huner, 2003). The imbalance in the PSII caused by cold stress might generate ROS, which can then damage the photosynthetic apparatus and the whole cell (Tyystjarvi, 2013). Therefore, tolerance to cold-induced photoinhibition can be considered a mechanism for cold tolerance. Additionally, RubisCO plays a central role in CO_2_ assimilation and photosynthesis efficiency. Crops that are acclimated to cold, such as winter wheat and rye, adjust their RubisCO content and are able to maintain a high CO2 assimilation rate (Yamori et al., 2010).

Low temperature can also affect the enzymes and ion channels responsible for maintaining the guard cell osmotic potential (Ilan et al., 1995). Due to the reduction in the photosynthetic capacity caused by cold stress, there is an increase in internal CO_2_ in the substotamal cavity, which reduces the stomatal aperture (Wilkinson et al., 2001).

In this study, GO enrichment analysis revealed enriched categories in up-regulated genes in GT1 related to photosynthesis compared to RRIM600 after 24 cold stress. Although we observed enriched GO categories related to plant defense in GT1 after 24h, we also identified enriched categories associated with photosystem II (GO:0009523), photosystem (GO:0009521), photosynthetic membrane (GO:0034357), chlorophyll biosynthetic process (GO:0015995), and photosynthesis (GO:0015979) (Supplementary Table 3). Additionally, the qPCR validation corroborated the *in silico* DEG analysis, in which the RubisCO gene was found to be up-regulated in GT1.

The pair-wise comparison between GT1 and RRIM600 revealed that the abscisic acid (ABA) receptor PYL4 gene was up-regulated in RRIM600 at 0 h, 90 minutes and 24 h of chilling stress. Additionally, after 24 h of cold stress, RRIM600 showed up-regulation of two abscisic acid receptor PYL4 genes and one ABA receptor PYR1 gene (Supplementary Table 3). The PYR/PYL ABA receptor genes are involved in ABA-mediated responses and play a major role in basal ABA signaling for vegetative and reproductive growth, modulation of the stomatal aperture and the transcriptional response to the hormone (Gonzalez-Guzman et al., 2012). Recent studies have demonstrated that overexpression of the PYR1 gene in poplar significantly reduces the content of H_2_O_2_ and significantly contributes to cold tolerance (Yu et al., 2017).

In GT1, the phototropin (PHOT1) gene, which is a blue-light receptor kinases that optimizes photosynthetic activity by sensing temperature and control the stomatal opening (Fujii et al., 2017, Sullivan et al., 2008), was up-regulated at 90 min and 12 h. We also observed that after 24 h of cold stress. GT1 showed up-regulation of a gene with high similarity to zeaxanthin epoxidase, which plays an important role in the xanthophyll cycle and alleviates the excitation pressure on the PSII reaction diverting photon energy into heat via zeaxanthin (Sui et al., 2007).

Furthermore, we identified one up-regulated gene in RRIM600 after 12 and 24 h of chilling stress with high similarity to the HT1 gene. In Arabidopsis, the HT1 serine/threonine kinase gene is reported to be involved in the control of stomatal movement in response to CO_2_. Additionally, genes with high similarity to hexokinase-1 and PtdIns3P 5-kinase (PI3P5K) were up-regulated in RRIM600 at 24 h of treatment. These genes are also reported to be related to stomatal closure via ABA (Bak et al., 2013) (Supplementary Table 2).

In Arabidopsis, SRK2E is a positive regulator in ABA-induced stomatal closure and is involved in stress adaptation. Additionally, SRK2E acts as a transcriptional repressor involved in the inhibition of plant growth under abiotic stress conditions (Yoshida et al., 2006). In this study, we identified the SRK2E gene as being up-regulated in RRIM600 after 90 minutes of cold treatment.

### 4.5 Molecular Markers for Rubber Tree Breeding

Molecular markers such as SSRs and SNPs are abundant in plant genomes. The development and genotyping of molecular markers is an important tool in genomic breeding and is the basis for genome selection (GS), genetic mapping and genome-wide association mapping (GWAS) as well as genetic linkage mapping. Next-generation sequencing allows the identification of thousands of putative SSR and SNP markers. Additionally, the identification of SNPs in genes using RNA-seq data allows the development of markers in candidate genes and the investigation of the variability of these genes in rubber trees (Mantello et al., 2014).

In this study, we identified a total of 27,111 putative SSRs and 264,341 putative SNPs. The putative molecular markers can be employed as a source to develop new markers for the species. The recent release of the *H. brasiliensis* genome, associated with the genome annotation, can be used to evaluate gene content and develop new markers in potential candidate genes. In this study, for example, we performed qPCR for a gene exhibiting high similarity to the ZAT10 gene. In Arabidopsis, ZAT10 plays an important role in cold tolerance and may be involved in the jasmonate early signaling response (Mittler et al., 2006). SNP calling revealed two SNPs in this gene in GT1.

Although a recent version of the genome was released containing 7,453 scaffolds, the rubber tree genome is 71% repetitive (Tang et al., 2016), which makes it difficult to assemble these scaffolds into chromosomal units. The development of new SSR markers can also be carried out and is an important tool helping link these scaffolds.

### 4.6 Identification of Alternative Splicing

RNA-seq experiments enable the detection of AS events. AS is a post-transcriptional modification of precursor mRNAs (pre-mRNAs) that can result in the formation of multiple distinct mRNAs from a single gene (Chamala et al., 2015). AS is an important mechanism for gene regulation in eukaryotes and can generate transcriptome and proteome diversity (Barbazuk et al., 2008; Chen and Manley, 2009). AS is involved in gene regulation that may regulate many physiological processes in plants, including the response to abiotic stresses such as cold stress (Chinnusamy et al., 2007; Tack et al., 2014).

In the rubber tree, the previous analyses of AS events were restricted to specific genes such as the rubber particle protein membrane (Chow et al., 2007) and the sucrose transporter genes (Dusotoit-Coucaud et al., 2009). Tang et al. (2010) evaluated the preferential expression of isoforms for the sucrose transporter. In their study, the authors found that the sucrose transporter isoform HbSUT3 was the predominant isoform expressed in rubber-containing cytoplasm (latex) (Tang et al., 2010).

A recent study in rubber tree using PacBio data identified AS in leaf tissues under normal conditions. Intron retention was the most abundant AS event identified, while exon skipping was less abundant, which agrees to the findings of the present study. Pootakham et al. (2017) identified 636 intron retention events corresponding to 41% of the total AS identified. However, these authors suggested that the number of identified AS is a small subset of the total possible number of AS events because their study used untreated leaf samples. In this study, we described AS events in rubber tree using leaf tissue under abiotic conditions for the first time. We detected a total of 9,226 intron retention events with a minimum of 10 reads supporting the AS sites, which corresponded to 45.5% of the total events identified. Alternative donor events represented a minority of the total number of detected AS events at 13%.

The results of in this study provide an overview of AS events in rubber tree based on non-stressed and cold stressed samples. Due to the importance of AS in plant adaptation, these data can be employed for further investigation of cold stress adaption in this species.

### 4.7 Rubber tree breeding

Compared to other crops, rubber tree domestication and breeding is recent, having started approximately 100 years ago (Mantello et al., 2014). Recent physiological studies involving rubber tree genotypes have demonstrated that RRIM600 is resistant to cold stress, while GT1 is cold tolerant, showing little leaf damage after 18 days of chilling exposure. It has been suggested that RRIM600 presents an “avoidance” strategy, in which it rapidly closes its stomata and down-regulates photosynthetic activity. Although GT1 is considered an intermediate tolerant genotype, this clone continues to grow and remains active with little leaf damage (Mai et al., 2010). The RNA-seq approach utilized in this study allowed a rapid and deep investigation of the genetic response under abiotic and biotic stresses in non-model species. The DEG analysis performed in this study provides insight to the genes up-regulated in each genotype, RRIM600 and GT1, across various durations of cold stress and corroborates the physiological findings of Mai et al. (2010). In this study, we observed that RRIM600 exhibits a more efficient ROS scavenging system compared with GT1, based on the large number of genes related to ROS scavenging that were up-regulated in RRIM600 during cold treatment. Moreover, important genes previously reported to be involved in stomatal closure were up-regulated in RRIM600. We also observed that genes related to cell growth were up-regulated in GT1 after 24 h hours of cold stress. Although we identified genes related to the defense response in GT1, the DEG and GO enrichment analyses showed that GT1 remains active, displaying up-regulation of genes related to photosynthesis during cold treatment. In addition, GT1 probably has a more efficient strategy for strengthening the cell wall, as revealed through GO enrichment analysis.

Rubber tree breeding programs are interested in genotypes resistant to cold and that exhibit latex production that does not cease during the cold season. The elucidation of different chilling tolerance strategies linked to information about possible genes involved in such responses, including the identification of molecular markers in these genes associated with information on AS events, provides a powerful tool for the genetic and genomic analysis of the rubber tree for breeding strategies and future studies involving GWAS and GS.

## Conflict of Interest

The authors have no conflicts of interest to declare.

## Author Contributions

Conceived and designed the experiments: CCM and APS. Performed the experiments: CCM and CCS. Analyzed the data: CCM, CS and LB. Contributed reagents/materials/analysis tools: CCM, LB, CCS, ESJ, PSG, BB, and APS. Wrote the paper: CCM. All authors read and approved the final manuscript.

## Funding

This work was supported by grants from the Fundação de Amparo à Pesquisa do Estado de São Paulo (FAPESP) (2007/50392-1;2012/50491-8), Conselho Nacional de Desenvolvimento Científico e Tecnológico (CNPq) (478701/2012-8;402954/2012), Coordenação de Aperfeiçoamento de Pessoal de Nível Superior (CAPES Computational Biology Program) and US National Science Foundation (IOS-1547787). Scholarships were provided by FAPESP to CCM (2014/18755-0) and CCS (2015/24346-9). APS and PSG are recipients of a research fellowship from CNPq.

## Acknowledgements

We would like to thank Dr. André Ricardo Oliveira Conson for helping with the sequences submission to NCBI database.

## Supplementary Material

**Supplementary Material 1. Rubber tree genome annotation.**

**Table 1. qPCR primer sequences of the 20 genes.**

**Table 2. Gene annotation of the DEGs identified for each time point in RRIM600 and GT1.**

**Table 3. GO enrichment of RRIM600 and GT1 at each time point series.**

